# Transformation of cereal grains: Botanical and chemical analysis of food residues encrusted on pottery from the Funnel Beaker settlement of Oldenburg LA 77, northern Germany

**DOI:** 10.1101/2023.12.11.571086

**Authors:** Lucy Kubiak-Martens, Tania F.M. Oudemans, Jan Piet Brozio, Dragana Filipović, Johannes Müller, Wiebke Kirleis

## Abstract

An integrated botanical and chemical approach is used to study surface residues on Funnel Beaker ceramics from the site of Oldenburg LA 77, in northern Germany. Organic residues were discovered adhering to fragments of thick-walled, undecorated ceramic vessels (n=19) and ceramic discs (n=2). The surface residues were studied using scanning electron microscopy (SEM), to examine remains of cereals and other plant tissues that survived food preparation and cooking, and using attenuated total reflectance Fourier transform infrared spectroscopy (ATR–FTIR) and direct time-resolved mass spectrometry (DTMS), to chemically identify specific food components. The SEM results show a reoccurring presence of cereal grain (emmer and barley) and one case of co-occurrence of emmer and fat-hen seeds. The SEM evidence for the use of sprouted emmer grain and milk-ripe barley from the Oldenburg residues greatly enhances our understanding of Neolithic foodways in northwestern Europe. The ATR-FTIR results showed that roughly a third of the surface residues contain traces of the original foods prepared or processed and DTMS results confirm that most of the residues primarily contain polysaccharides and a minimal amount of plant protein and that they lack lipids. Only one residue presents minor indications for a (partly) animal origin. The ceramic vessels were thus used almost exclusively for the processing or cooking of cereal grains. This study offers an intimate view of the cuisine and cooking practices (and in some cases their seasonal timing) in an early agricultural village located in a marginal farming region on the south coast of the Baltic Sea.

## Introduction

The subsistence economy of the first farmers in northern Europe, the Funnel Beaker groups, has been intensively studied during the past two decades [1-14]. It is well known, which cultivated plants and domestic animals were used and regional variations have been identified. Although the study of stable isotopes has provided knowledge about cultivation practices [10-12], animal mobility [13], and food pathways [14], less is known about what items found their way into cooking vessels and how daily meals were prepared. The first efforts to look into cooking vessels with Baltic provenience were undertaken on sherds from the Mesolithic-Neolithic (Ertebølle–Funnel Beaker) transitional sites of Neustadt, in Holstein, and Aakonge and Stenø, in Denmark [15-17].

Recent developments in the application of high-resolution microscopy applied in archaeobotany have brought additional advances in analysis of the actual remains of processed food or cooked meals from charred objects and food residues encrusted on the surface of ceramic vessels [18-21]. These applications, especially when combined with the chemical analysis of surface residues, redress the imbalance that resulted from the method-driven focus in biomarker research on animal lipids, to reveal the importance of the plant component in ancient cuisine.

In this study, surface residues are analysed using an integrated botanical and chemical approach to understand how meals were prepared in the cooking vessels recovered from the Middle Neolithic settlement of Oldenburg LA 77, which is one of several Neolithic sites located in the Oldenburger Graben microregion in northeastern Schleswig-Holstein, Germany (Fig 1). The site is attributed to the Funnel Beaker (German: Trichterbecher, or TRB) groups and dates to 3270–2920 cal. BCE. With the agglomeration of people in c. 40 houses at the peak of its occupation, it is one of the oldest villages in Schleswig-Holstein and illustrates the transformation of social life in the Middle Neolithic, from single farmsteads to a village community. Oldenburg LA 77 was a fully agrarian settlement, with a typical Funnel Beaker economy, comprising distinctive cultivation practices for cereals, such as emmer (*Triticum dicoccum*) and naked barley (*Hordeum vulgare* var. *nudum*); the tending of livestock; hunting; fishing; and the gathering of wild plants [1, 6, 10].

**Fig 1.**
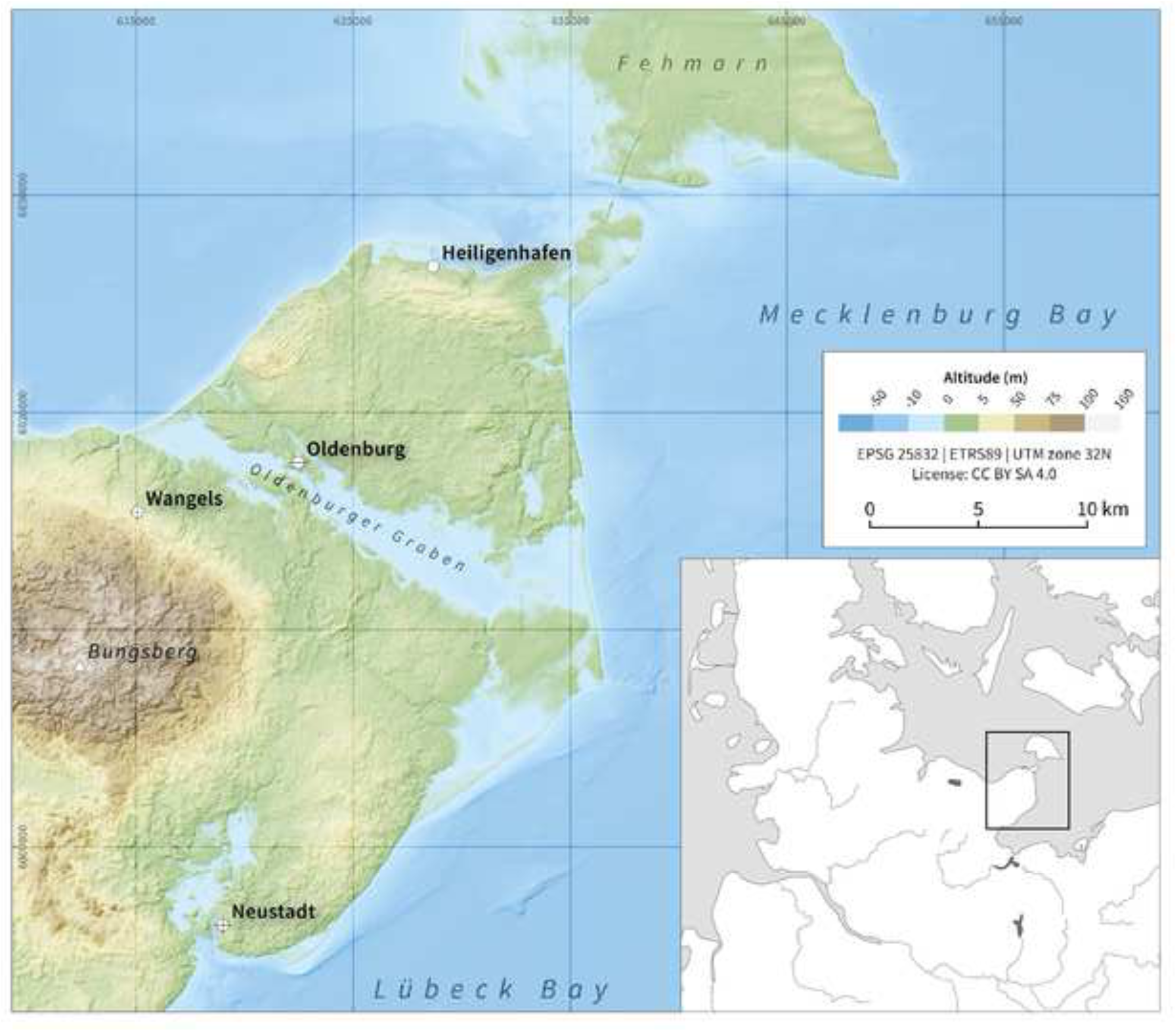
Map showing the location of Oldenburg LA 77 within the Oldenburger Graben area. Figure: J.P. Brozio. Edited by D. Filipović from [24] under the CC BY license 4.0, with permission from the authors.

An earlier study of absorbed lipids that were preserved inside the ceramic matrix of a number of sherds from Oldenburg LA 77 gave a first insight into cooking practices at Oldenburg LA 77 [22]. Weber and colleagues combined biomarker analysis and isotope ratio information to conclude that dairy products and plant oils had been prepared in the vessels studied. This paper focuses on residues encrusted on the surface of ceramic vessels because they can provide more detailed and more direct information about the vessel use than the study of lipids absorbed into the ceramic matrix which tends to show an overrepresentation of animal fat and often miss the plant element. The first benefit of surface crusts over absorbed lipids is that surface residues represent the last, or one of the last, use-phases of the vessel, whereas absorbed lipids represent the cumulative deposition over the entire use-life of a vessel. The second benefit is that surface residues can be studied for a full range of food components (e.g., proteins, lipids, and polysaccharides such as starches and sugars), as well as for any remaining small fragments of plant and or animal tissue, thus yielding much more detailed information about food sources. The third benefit is that the range of detectable food components is not limited to those containing lipids, making many plant sources more visible.

In the case of crops, food preparation practices represent the culmination of the long and laborious production process that transforms the sown grain into a desired meal. This transformation is largely elusive in the archaeological record and is usually reconstructed based on indirect evidence, such as food processing tools. Studying food crusts can provide valuable information about the original vessel contents. Because they are found firmly attached to either the interior or the exterior surface of ceramics, they form a direct source of information about food choices, the preparation and processing of food before cooking, the mixing of foods, and even the use of different types of pots for similar or, indeed, different purposes.

The application of an integrated botanical and chemical approach to surface residue studies has been proven to render a more complete understanding of the cooking practices and food sources used in prehistoric times [20-21, 23]. The use of scanning electron microscopy (SEM) enables the observation and identification of various plant particles, and some animal remains that survived the processes of food preparation and cooking. The high-resolution microscopic analysis also enables the taxonomic identification of these remains. The plant particles are often minute fragments of grain tissue (e.g., pericarp or aleurone); cereal light chaff (including fragments of palleas, lemmas, and glumes); remains of seeds; and vegetative plant parts, whereas the animal remains are often small bone fragments or fish scales. The chemical approach combines two types of analysis: attenuated total reflection– Fourier-transform infrared spectroscopy (ATR–FTIR), for a first general identification of the nature of the residues and their state of preservation, with direct time-resolved mass spectrometry (DTMS), a technique that makes it possible to further characterise the entire organic composition of the charred residue without prior treatment, including volatile, extractable compounds and solid, non-extractable compounds.

The detection of any organic amorphous remains - that cannot be analysed by other means - helps identify the natural origin of the surviving residues and explain their presence in terms of human behavior in the past.

## The site and its economy

### The Funnel Beaker settlement site of Oldenburg LA 77

The settlement site of Oldenburg LA 77, dated to 3270–2920 cal BCE, is situated on a dune-like peninsula of ca. 3 ha in the western part of a low-lying wetland area on the southern Baltic coast, the so-called Oldenburger Graben (Fig 1). Initially a fjord, the area transformed into a lagoon at the turn of the 4th to the 3rd millennium BCE [1, 24-26]. The settlement was part of a larger settlement system that used (former) islands and peninsulae as residential areas and land along the boundaries of the wetlands as agricultural land. Associated burial sites are located on the adjacent moraine ridges to the south and north of the Oldenburger Graben [1-2, 24-27].

The archaeological features in the excavated parts of the site indicate the presence of five ‘houses’ and several ‘huts’ [1]. Although the houses cluster on higher terrain, remains of domestic origin in various states of preservation have been found in a cultural layer extending into the wetlands at the south shore of the Neolithic settlement. Some of these artefacts and ecofacts are secondary deposits. Extensive ^14^C AMS dating of the site, in combination with the seriation of ceramics following the domestic pottery typology in the southern range of the Funnel Beaker North Group for the period 3270–2920 cal BCE [1, 27], distinguished four occupation phases: Oldenburg 1: 3270–3110 cal BCE; Oldenburg 2: 3110–3020 cal BCE; Oldenburg 3A: 3020–2990 cal BCE; and Oldenburg 3B: 2990–2920 cal BCE.

### Evidence of plant use at the Funnel Beaker settlement site of Oldenburg LA 77

Previous insights into plant use at Oldenburg LA 77 came from two classes of archaeological materials: macro-and microbotanical remains. Trenches 1, 8, and 10 (Fig 2) yielded large amounts of well-preserved macroscopic plant remains, and these have been studied in detail [1, 6]. The vast majority of this material was charred or partially charred. Uncharred (in this case waterlogged) plant remains were primarily found in the cultural deposits in the wetland area at the southern edge of the peninsula.

**Fig 2.**
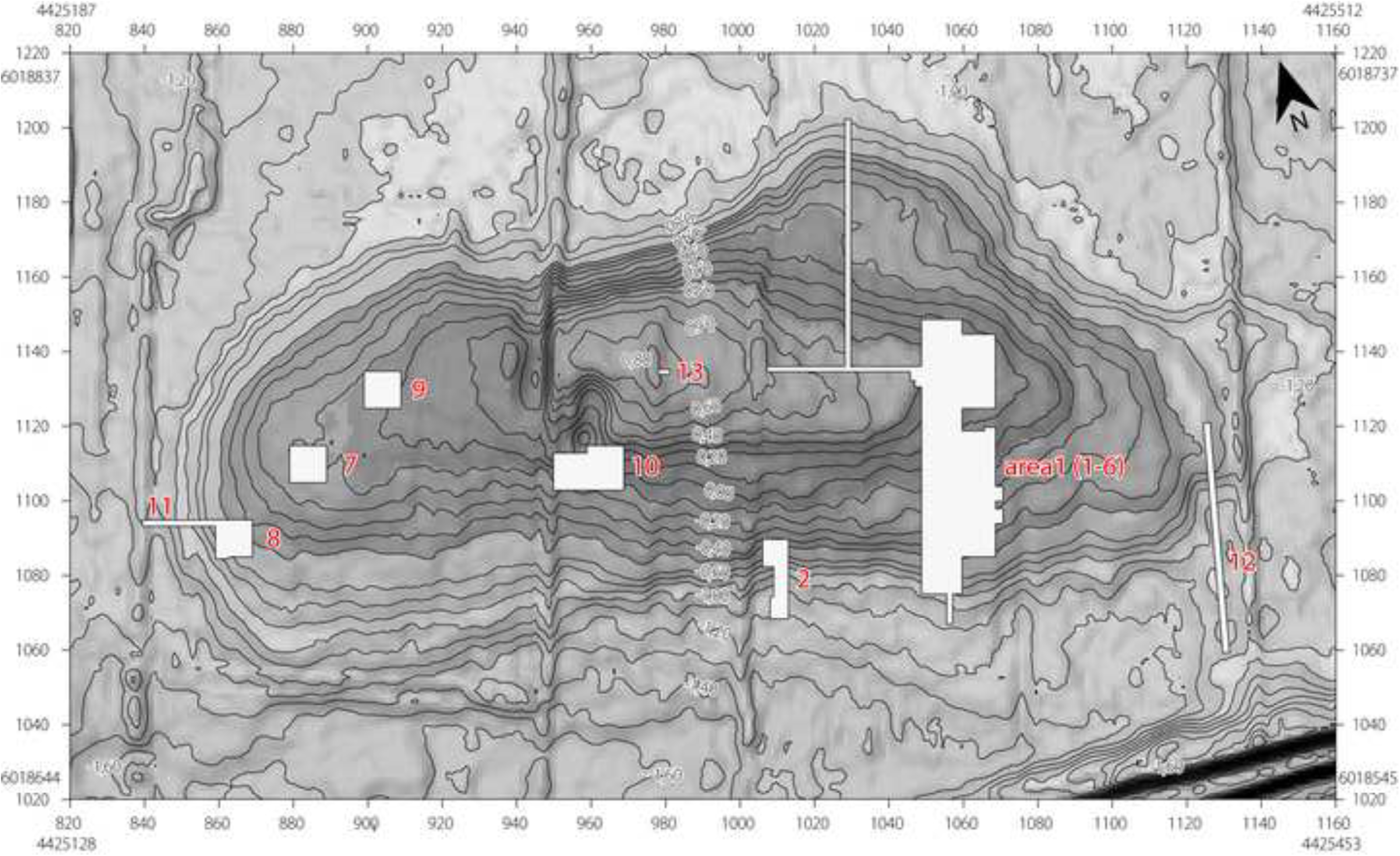
The reconstructed settlement area (dark grey shading) of Oldenburg LA 77, showing the location of the trenches excavated in 2009–2012 (white rectangular areas). Trenches 1 and 2 extended into the wetland area (light grey shading) at the southern edge of the peninsula. Figure: K. Winter; map base: DGM ©LVermGeo SH. Reprinted from [1, Abb. 13], with permission from the copyright holders; original © 2016 by UFG CAU Kiel and author.

Archaeobotanical analysis revealed the presence of cultivated and wild taxa that are commonly part of the plant assemblage from Funnel Beaker sites. The crop plants primarily consist of two cereals, namely naked barley and emmer. Einkorn (*T. monococcum*) and free-threshing wheat (*T. aestivum/durum/turgidum*) are present in minute amounts. Stable isotopic analysis on some of the charred grains revealed distinct cultivation strategies for the different cereals [10]. Single seeds of flax (*Linum usitatissimum*) and opium poppy (*Papaver somniferum*) were also identified, as were remains of edible wild plants, not only nuts and fruits from common gathered plants, such as crab apple and hazel but also fat-hen, small-seeded grasses and sedges [1, 4, 6, 10].

### Evidence of animal use at the Funnel Beaker settlement site of Oldenburg LA77

Zooarchaeological analysis has so far revealed a dominance of the remains of domestic animals, with cattle and pig more frequent than sheep, goat and dog. The remains of wild animals show the presence of large and small mammals (deer, elk, hare), fish and birds [1, 2]. The ratios of domestic to wild animals, as well as the ratios among the varied species of domestic animals, remain constant throughout the duration of the settlement, pointing to continuity in animal husbandry. The ratios of individual species of wild animals, however, vary slightly among different phases of occupation, possibly indicating changes in their availability. The presence of remains of large animals – cattle, deer, elk – suggests that the butchering and consumption were practiced on-site and perhaps by a larger group of people [1, 2].

## Materials and methods

### Potsherds with food crusts

After excavation, the ceramics from the site had been air-dried and then placed in plastic bags. Following visual inspection of the pottery assemblage, 21 pottery fragments with residues encrusted on their surface were selected for the present study (Table 1 and S1). The sherds come mainly from undecorated vessels with a conical top (OLD 03, 16 and 21) or cylindrical rim (OLD 02, 07 and 10). All vessel sherds originate from large vessels, with thick walls, made of coarse to very coarse clay mixed with crushed granite. The range of colours suggests different firing conditions (possibly including secondary burning) and different clay types. Two fragments (OLD 08 and 15) were originally part of ceramic discs (*Tonscheiben)* and have decoration in the form of a row of fingertip and fingernail impressions along their edges (S1). The function of these ceramic discs has not been resolved, even though they are regularly recovered from prehistoric sites in Europe [27]. One suggested use for these ceramic discs is the baking of flatbread; hence they are also referred to in German as *Backteller* (baking plates). They are also thought to have served as lids and serving plates [27-28].

**Table 1.** Details of Oldenburg LA 77 ceramic fragments from which samples of surface residue were taken for botanical and chemical analyses. frag. = fragment; undecor. = undecorated

| Sample Identifier | ALSH Inventory Number | Find Number | Find Context | Type of Ceramic Object | Sampling Location | Residue Description | SEM | FTIR | DTMS | Date Range | Fig |
| --- | --- | --- | --- | --- | --- | --- | --- | --- | --- | --- | --- |
| OLD 01 | 2010-663.3082 | 232 | Submerged zone, cultural detritus | Body frag. of undecor. vessel | Interior wall | Brown-black residue, medium layer, 1–2 mm thick, multiple patches subsampled | yes | yes | yes | 3270–2920 cal. BCE | . |
| OLD 02 | 2010-663.3087 | 309 | Submerged zone, cultural detritus | Cylindrical rim frag. of undecor. vessel | Exterior rim | Black, thin residue, thin/medium layer, <1 mm thick | yes | yes | yes | 3270–2920 cal. BCE | . |
| OLD 03 | 2010-663.3096 | 423 | Submerged zone, cultural detritus | Conical top frag. of undecorated vessel | Exterior rim | Black residue, medium/thick layer, 1–2 mm thick | yes | yes | no | 3270–2920 cal. BCE | . |
| OLD 04 | 2010-633.3099 | 154 | Submerged zone, cultural detritus | Body frag. of undecor. vessel | Interior wall | Black residue, medium layer, 1–2 mm thick | yes | yes | no | 3340–2937 cal. BCE (AMS-date on food crust) | 4 |
| OLD 05 | 2011-432.3093 | 382 | Submerged zone, cultural detritus | Body frag. of undecor. vessel | Exterior rim | Black residue, 1–2 mm thick layer | yes | yes | no | 3270–2920 cal. BCE | 7 |
| OLD 06 | 2010-633.3101 | 546 | Submerged zone, cultural detritus | Body frag. of undecor. vessel | Interior wall | Black residue, medium layer, 1–2 mm thick | yes | yes | yes | 3321–2914 cal. BCE (AMS-date on food crust) | . |
| OLD 07 | 2010-633.3094 | 197 | Submerged zone, cultural detritus | Cylindrical rim frag. of undecor. vessel | Interior rim | Thin brown and white spots, thin/medium layer, <1 mm thick | yes | yes | no | 3328–2098 cal BCE; median 3067 cal BCE | . |
| OLD 08 | 2011-432.7915 | 2001 | Submerged zone, cultural detritus | Ceramic disk (*Tonscheibe*) | On top of disc | Thin/fine black residue, <1 mm thick (sherd sampled earlier for absorbed residue analysis) | no | yes | yes | 3270–2920 cal. BCE | . |
| OLD 09 | 2011-432.7907 | 2158 | Submerged zone, cultural detritus | Body frag. of undecor. vessel | Interior wall | Brown-black residue, thick layer, 3 mm | yes | yes | yes | 3270–2920 cal. BCE | 12 |
| OLD 10 | 2011-432.8108 | Ge-112 | Pit | Cylindrical rim frag. of undecor. vessel | Interior rim and wall | Brown-black residue, thick layer, 3–4 mm | yes | yes | no | 3270–2920 cal. BCE | 16 |
| OLD 11 | 2011-432.8096 | 2109-01 | Submerged zone, cultural detritus | Body frag. of undecor. vessel | Interior wall | Brown-black residue, thick layer, 3–4 mm | no | yes | yes | 3337–2935 cal. BCE (AMS-date on food crust) | . |
| OLD 12 | 2011-432.8096 | 2109-02 | Submerged zone, cultural detritus | Body frag. of undecor. vessel | Interior wall | Brown-black residue, thick layer, 3–4 mm | yes | yes | no | 3270–2920 cal. BCE | . |
| OLD 13 | 2010-633.3073 | 5007 | Posthole | Body frag. of undecor. vessel | Interior wall | Black residue, thin layer, 1 mm | no | yes | no |  | . |
| OLD 14 | 2010-633.3083 | 147 | Occupation horizon/layer | Body frag. of undecor. vessel | Exterior wall | Brown-black residue, medium layer, 1–2 mm thick | no | yes | no | 3270–2920 cal. BCE | . |
| OLD 15 | 2010-633.3092 | 308 | Submerged zone, cultural detritus | Ceramic disk (*Tonscheibe*) | On top of disc | Brown and white residue, less than 1 mm thick (sherd sampled earlier for absorbed residue analysis) | no | yes | yes | 3270–2920 cal. BCE | . |
| OLD 16 | 2010-633.3100 | 334 | Submerged zone, cultural detritus | Undecor. vessel with conical top | Exterior rim | Thin brown-black residue, shiny, <1 mm thick | no | yes | no | 3270–2920 cal. BCE | . |
| OLD 17 | 2010-633.3102 | 95 | Occupation horizon/layer | Body frag. of undecor. vessel | Interior base | Light brown residue, <1 mm thick (sherd sampled earlier for absorbed residue analysis) | no | yes | no | 3270–2920 cal. BCE | . |
| OLD 18 | 2011-432.7908 | 2361 | Submerged zone, cultural detritus | Body frag. of undecor. vessel | Interior wall | Thin brown-black residue, < 1 mm thick | no | yes | no | 3270–2920 cal. BCE | . |
| OLD 19 | 2011-432.7908 | 2361 | Submerged zone, cultural detritus | Body frag. of undecor. vessel | Exterior wall | Black residue, medium layer, 2 mm thick | no | yes | no | 3270–2920 cal. BCE | . |
| OLD 20 | 2011-432.7910 | 2440 | Submerged zone, cultural detritus | Body frag. of undecor. vessel | Interior wall | Brown-black residue, 2 mm thick | no | yes | no | 3270–2920 cal. BCE | . |
| OLD 21 | 2011-432.7912 | 2088 | Submerged zone, cultural detritus | Conical top frag. of undecorated vessel | Interior rim | Thin brown residue, <1 mm thick | no | yes | no | 3270–2920 cal. BCE | . |

Most of the analysed potsherds (OLD 01–07, 09, 11, 12, 15, 16 and 18–20) come from the deposits in the southernmost parts of trenches 1 and 2 (Fig 3), the formerly submerged zone where cultural detritus accumulated during the entire timespan of the settlement, that is, ca. 3270–2920 BCE (there are also traces of younger depositions in the colluvium). It is, therefore, difficult to associate these fragments with a specific settlement phase. The two ceramic disc fragments (OLD 08 and 15) can be typo-chronologically assigned to the Nordic Middle Neolithic I–V, that is, 3300–2800 BCE. Two of the vessel sherds (OLD 14 and 17) were found in the use-horizon and another (OLD 13) was found in a posthole; neither context can be dated more precisely than the general period of settlement occupation, that is, 3270– 2920 cal BCE. Sherd OLD 10 was discovered in a pit assigned to a group of features most probably originating from the Oldenburg 2/3A phase, that is, 3110–2990 cal BCE.

**Fig 3.**
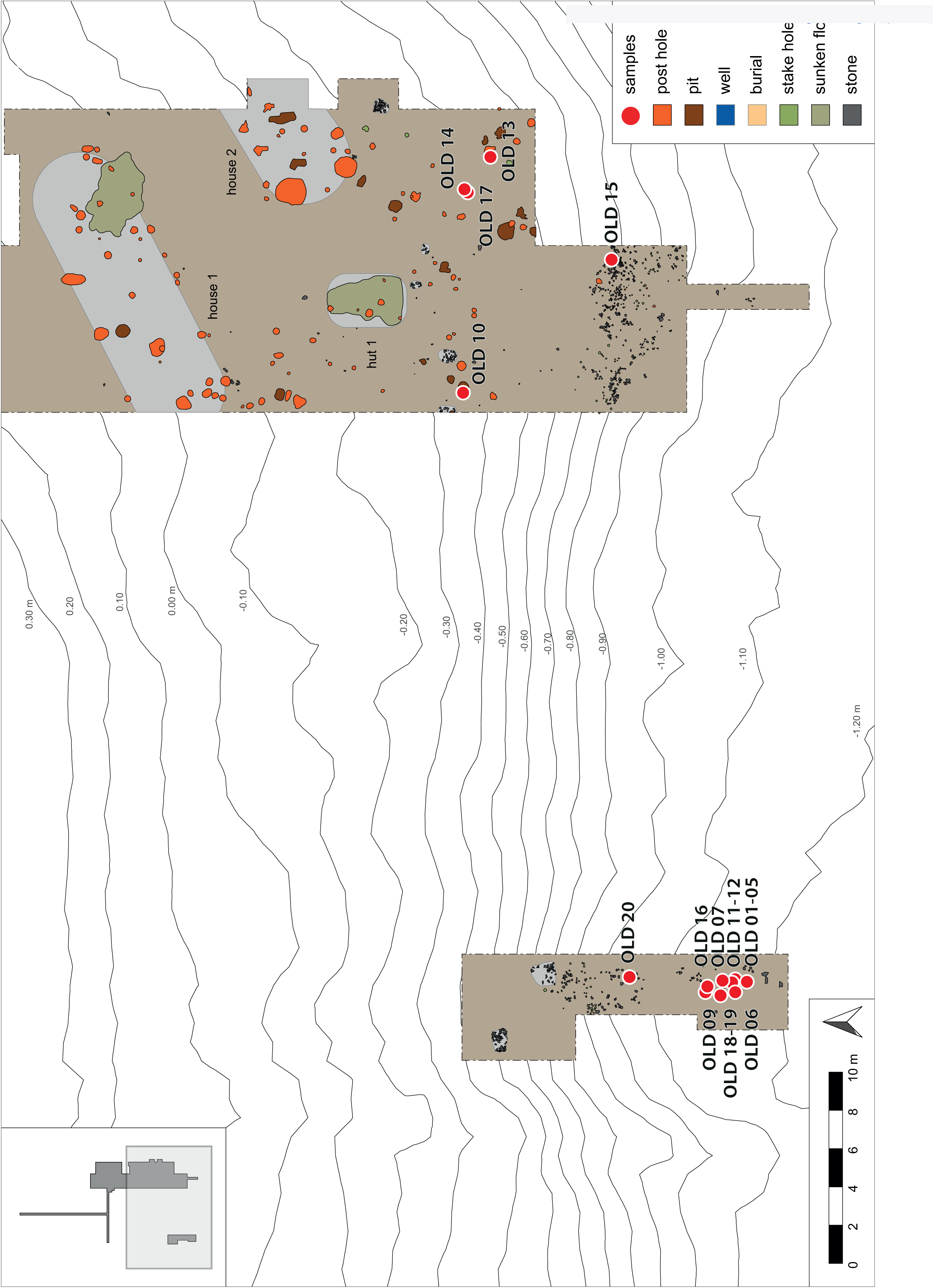
Find location of the sampled potsherds in trenches 1 and 2 at Oldenburg LA 77. All sherds originate from the wetland area at the southern edge of the settlement. Most come from a cultural layer, possibly as secondary deposits. One sherd originates from a posthole (OLD 13) and one from a settlement pit (OLD 10). Modified by J.P. Brozio from [1, Abb. 13], with permission from the copyright holders; original © 2016 by UFG CAU Kiel and author.

Subsamples of food crust removed from three of the potsherds recovered from the colluvium (OLD 04, 06 and 11) were submitted for radiocarbon dating. The results, as well as those for two emmer grains recovered from the occupation layer within trench 1, are presented in S2.

### Sample selection, pre-treatment and analyses

The ceramic assemblage was first examined with the naked eye to identify sherds with adhering residues. This visual scan yielded 21 sherds, with residues on the interior (n=13) or exterior surface (n=6) of the sherds or on the top surface (i.e. the face of sherd with fewer signs of fire exposure) of the ceramic discs (n=2). The surface residues on these 21 ceramic fragments were then examined using a Leica microscope with incident light and a magnification range of 6–50× to describe the macroscopic characteristics of the residues and to determine the potential for further analyses. Some of the residues displayed exceptionally good preservation (see examples in Figs 4 and 7). The residues varied in thickness from thin (c. 1 mm or less) to medium (c. 2 mm) and (in most cases) thick (2–3 mm); in colour from black to brown; and texture from solid to porous or spongy (see Table 1).

**Fig 4.**
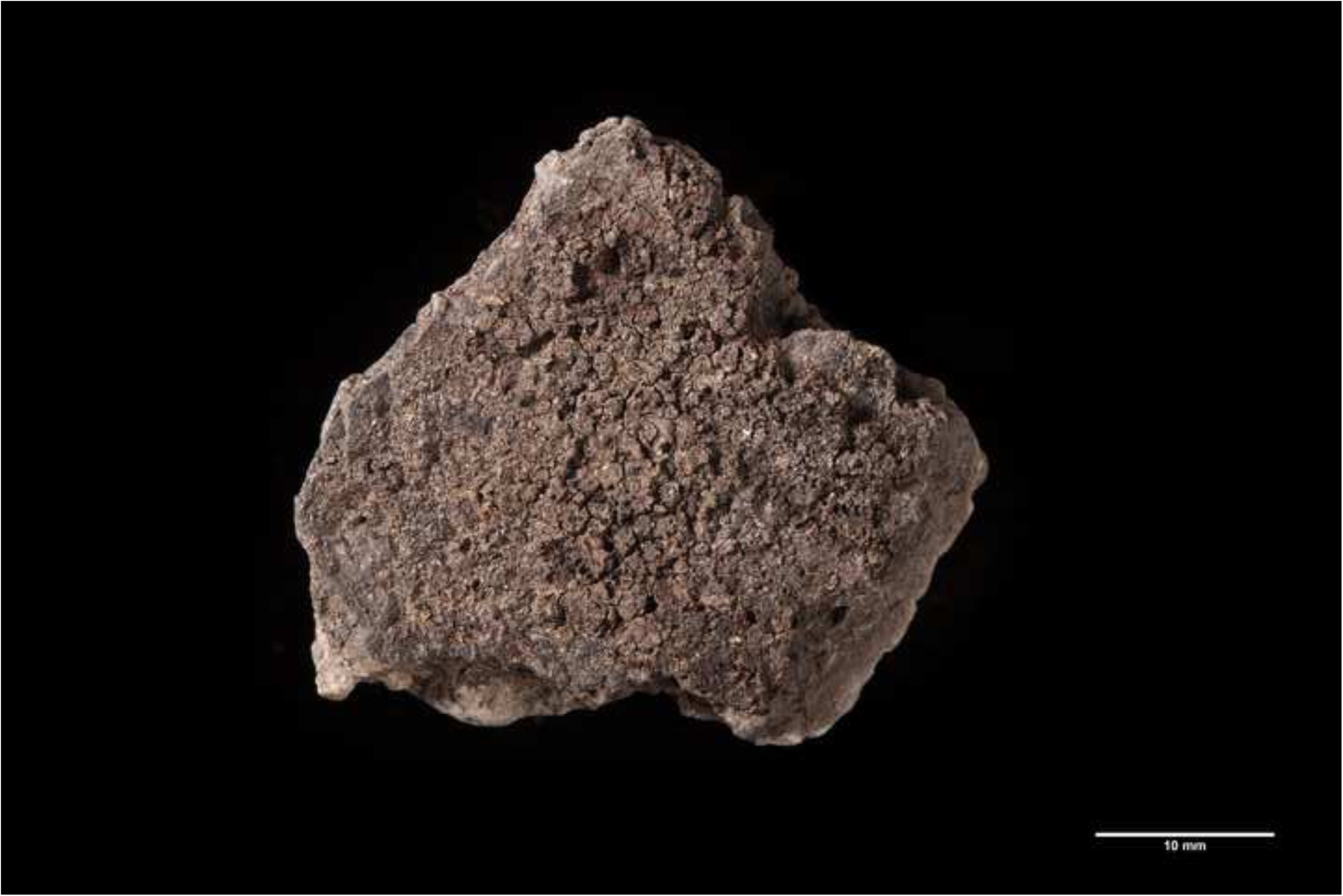
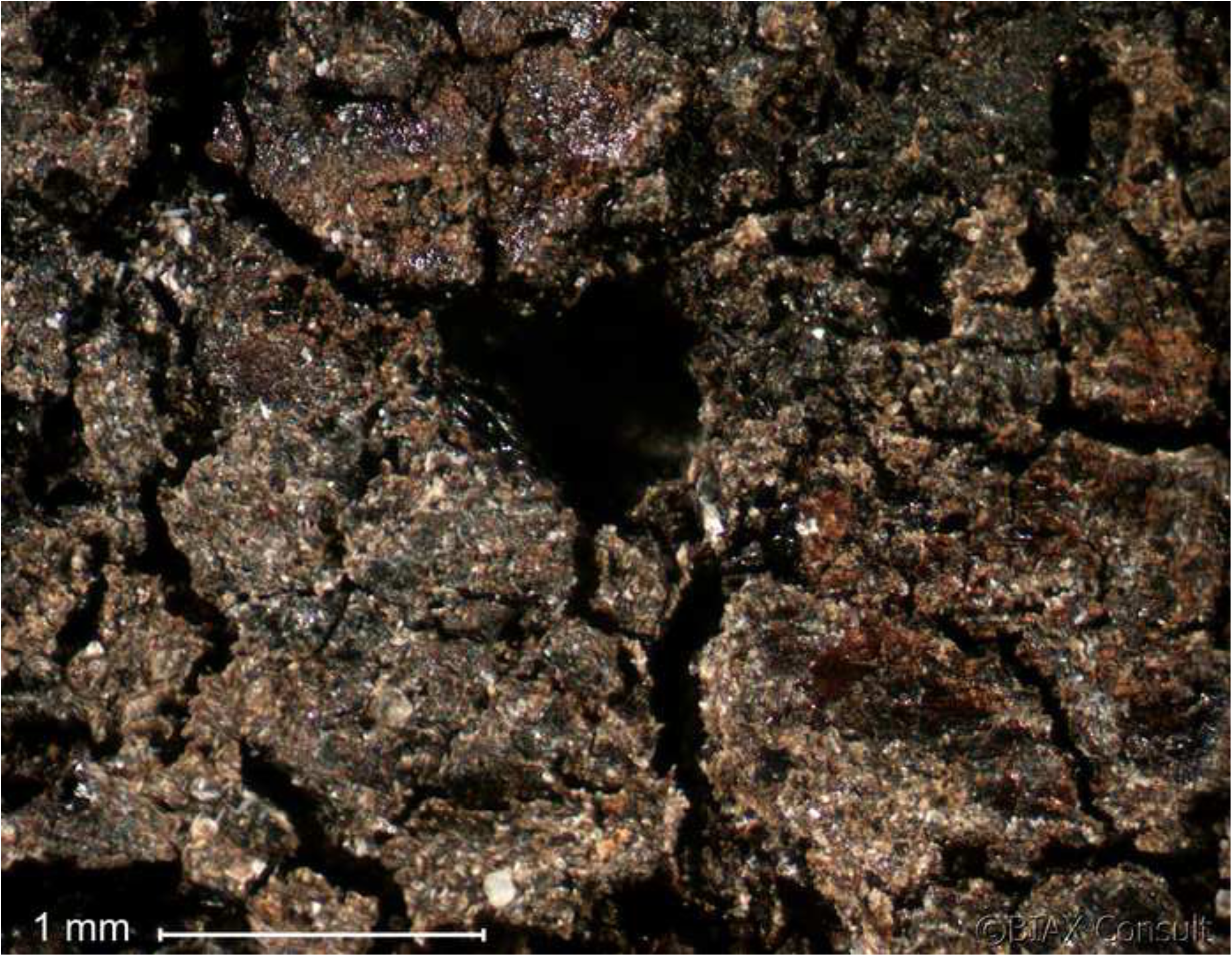
Oldenburg LA 77, sherd OLD 04, binocular microscope images of the surface of the food crust, showing the upper face of the surface residue (left) and detail of the area indicated by the white square in the left photo, showing grain particles embedded in the residue matrix (right). Images: BIAX/W. van der Meer.

SEM analysis was performed on the residues from 10 of the 21 sherds (see Table 1) using the JEOL JSM-6480L instrument at the Naturalis Biodiversity Center in Leiden, the Netherlands. Multiple subsamples were taken from various parts of each surface residue, to ensure wide coverage and account for potential variation in composition within the crust.

Portions of the residue were detached from each sherd and mounted on the SEM stubs using double-sided carbon tape strips. The samples were sputter-coated with a layer of platinum–palladium approximately 20 μm thick. The SEM images were captured at 30–750× magnification. The analysis centered mainly on the lower surface of the residues, i.e., the surface that had been attached to the pot, and on their interior; in several cases, the upper surface of the residues was also examined.

Chemical analysis was performed on the residues from all 21 sherds (see Table 1). A combination of ATR–FTIR and DTMS was applied. FTIR is a rapid analytical technique that is useful for the initial screening of solid samples, without necessitating further preparation. DTMS is a more powerful technique, employed for detailed analysis of very small samples of complex solid organic materials. DTMS makes it possible to characterise the complete organic composition of the material, including volatile, extractable compounds and solid, non-extractable compounds. DTMS results provide information about a broad range of compounds, such as lipids; waxes; terpenoids; saccharides; small peptides and protein fragments; and a broad range of thermally stable, more or less condensed polymeric components (commonly called ‘chars’ or ‘carbonised material’).

Best practice involves taking samples after the removal of the outer surface of the residue, in order to limit impregnation with soil material or possible post-excavation contamination. However, due to the limited amount of residue present, it was not feasible to remove the outer surface of all of the residues before sampling (see Table 1).

Instrumental conditions were as follows. In preparation for the FTIR measurements, ca. 10 µg from each sample was applied to the diamond window and flattened using the pressure arm of the ATR unit. FTIR analysis was performed in the ATR modus of the Fourier-transform infrared spectrometer (Thermo Scientific Nicolet iS05 equipped with an iD7 ATR unit). The spectral resolution was 4 cm^-1^ and the spectral range was 4000–550 cm^-1^. A total of 64 scans were collected per measurement, and the scans were analysed using the OMNIS software. The spectra were transformed using the functions “ATR correct”, “Automatic baseline correct” and “Transmission” to result in the depicted spectra. Prior to the DTMS analysis, ca. 50 µg of each sample was suspended in 5 µL of DCM/methanol (1:1 vol.) and homogenised. A small amount of the suspension was applied to the filament of the mass spectrometer, dried and subsequently analysed. DTMS was performed using a ISQ 7000 Thermo Scientific single quadrupole mass spectrometer equipped with a Direct Exposure Probe (DEP) and combined with a Thermo Scientific direct probe controller. Source temperature was 250 °C, the temperature programmed initial current was 0 mA (5 seconds) ramping to 1000 mA (with 10 mA/s) and held at 1000 mA for an additional 20 seconds. The following MS conditions were applied: the electron ionisation energy was 18eV, the scanning range was *m/z* 45–100 and the scanning speed was 5 scans per second. The data were collected and processed using the Xcalibur software.

## Results

The results on the analyses of the organic residues adhering to potsherds from undecorated Funnel Beaker ceramic vessels from Oldenburg LA 77 are presented starting with the SEM analysis, followed by the FTIR and DTMS analyses.

## SEM analysis

### Food crusts with emmer

The SEM analysis revealed the presence of emmer grain particles in five residues (OLD 01, 04, 06, 11 and 12 as illustrated in Figs 5, 6 and 8), one of which also contained fragments of fat-hen seeds (OLD 06 as illustrated in Fig 9). The most frequently observed grain tissue was single-layered aleurone tissue. In cereal grains, aleurone is the outermost layer of the starchy endosperm, and it stores the main protein content [29]. It is noteworthy that single-layered aleurone is present in all of the cultivated cereals except for barley, which develops multiple aleurone layers [30-31]. In the food crusts from Oldenburg LA 77, the single-layered aleurone was often accompanied by pericarp tissue. The characteristic pattern of the transverse cells, in some cases accompanied by longitudinal cells (Fig 8 left), and their association with the single-layered aleurone, narrowed the taxonomic identity down to *Triticum*, most likely emmer grain.

**Fig 5.**
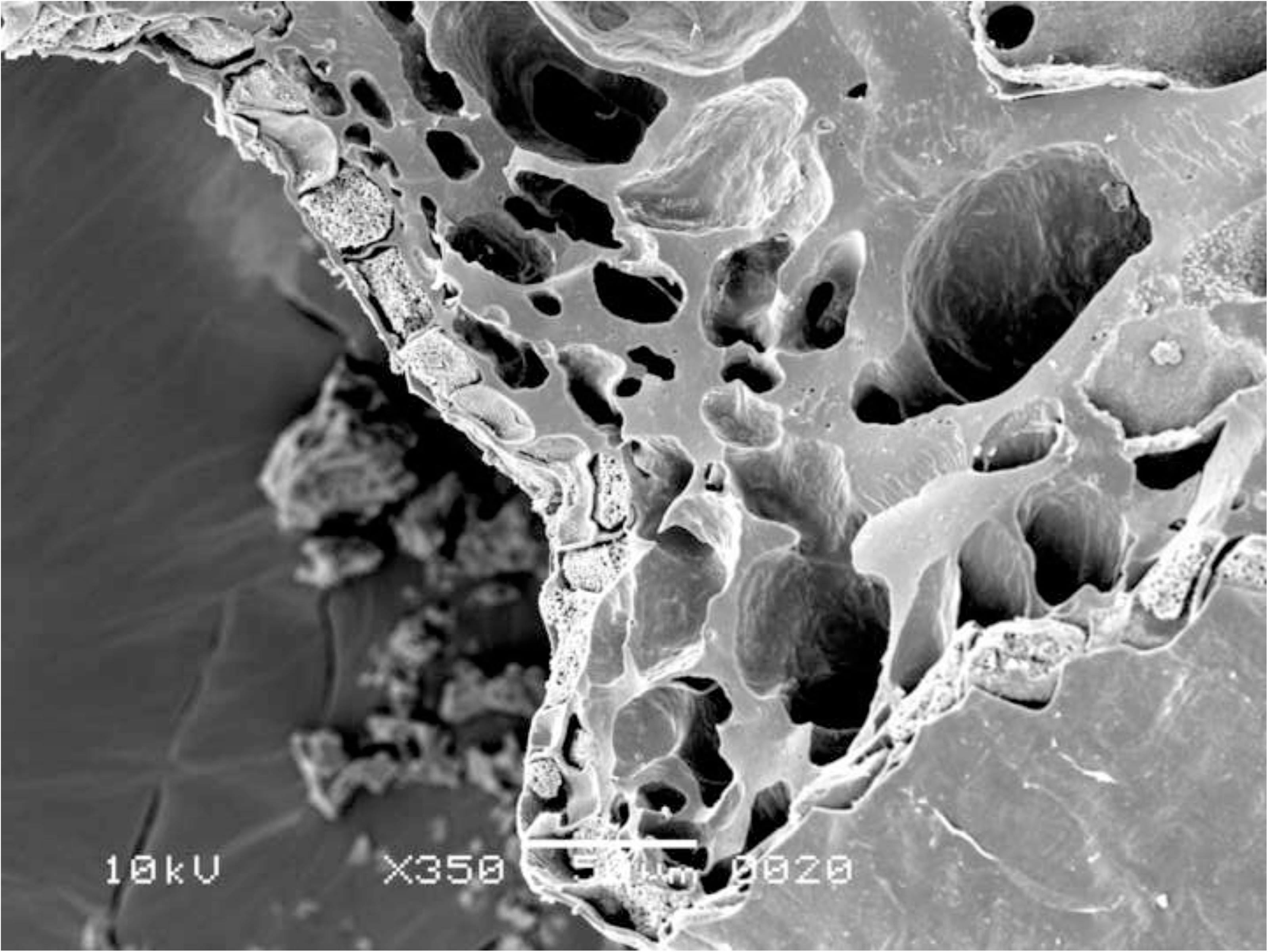
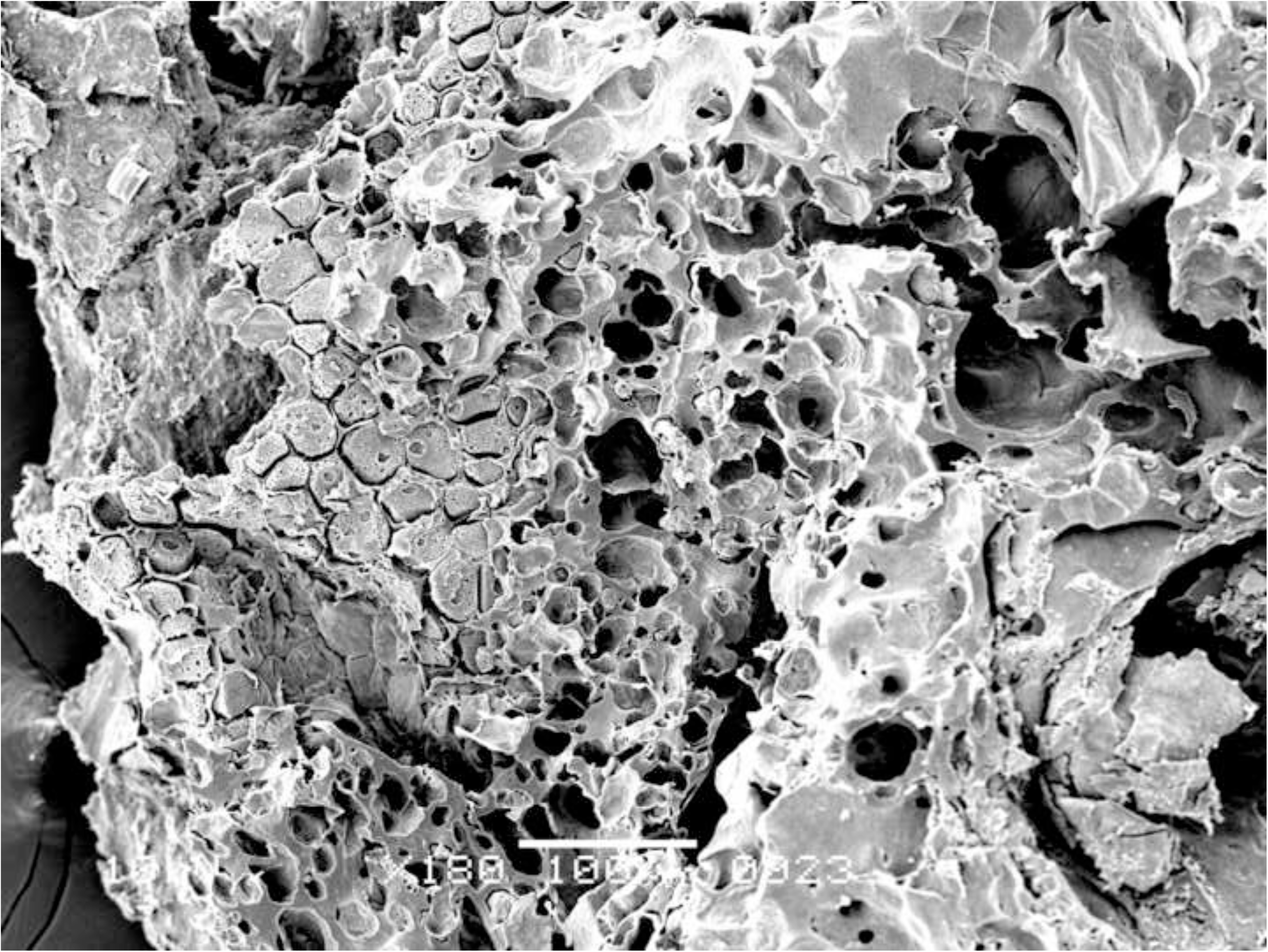
Oldenburg LA 77, sherd OLD 04, SEM images of the internal microstructure of the food crust, showing grain fragments with single-layered aleurone and starchy fused grain endosperm (in transverse view) (left) and single-layered aleurone in top view (right), both indicating *Triticum* (here most likely emmer). Images: L. Kubiak-Martens.

**Fig 6.**
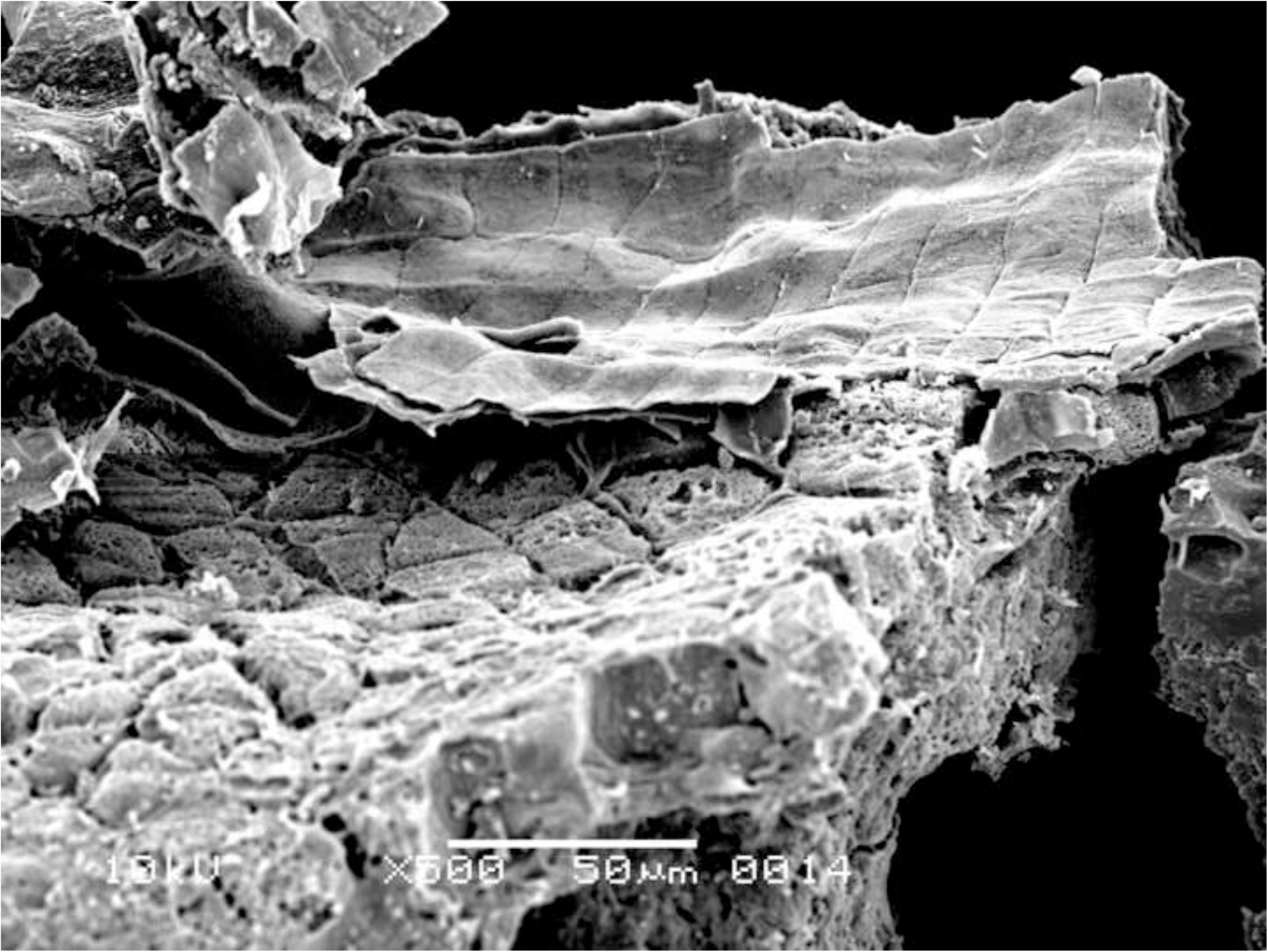
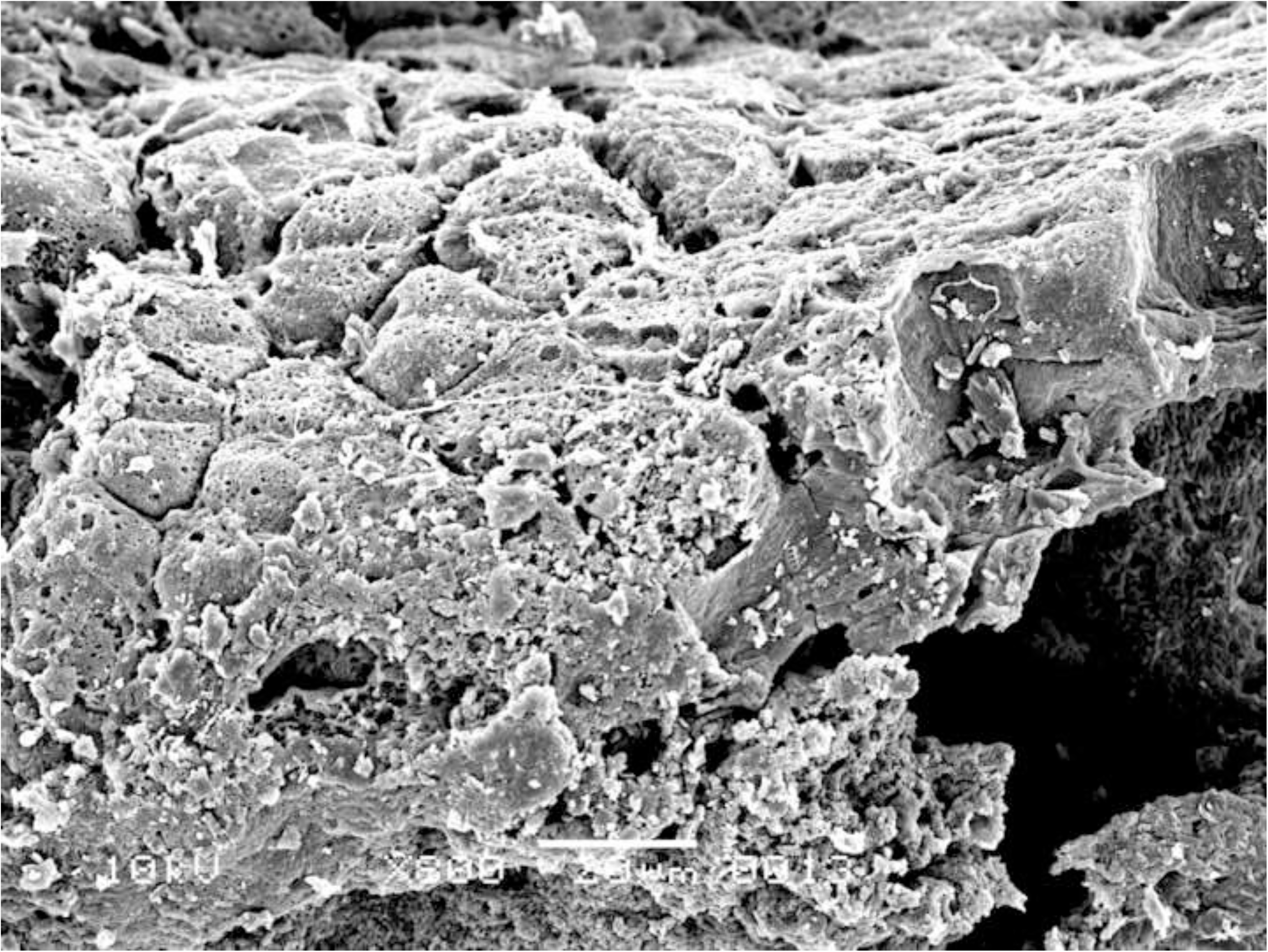
Oldenburg LA 77, sherd OLD 01, SEM images of the internal microstructure of the food crust, showing single-layered aleurone and pericarp tissue (left) and single-layered aleurone separated from the other grain tissues (right), indicating *Triticum* (here most likely emmer). Images: L. Kubiak-Martens.

**Fig 7.**
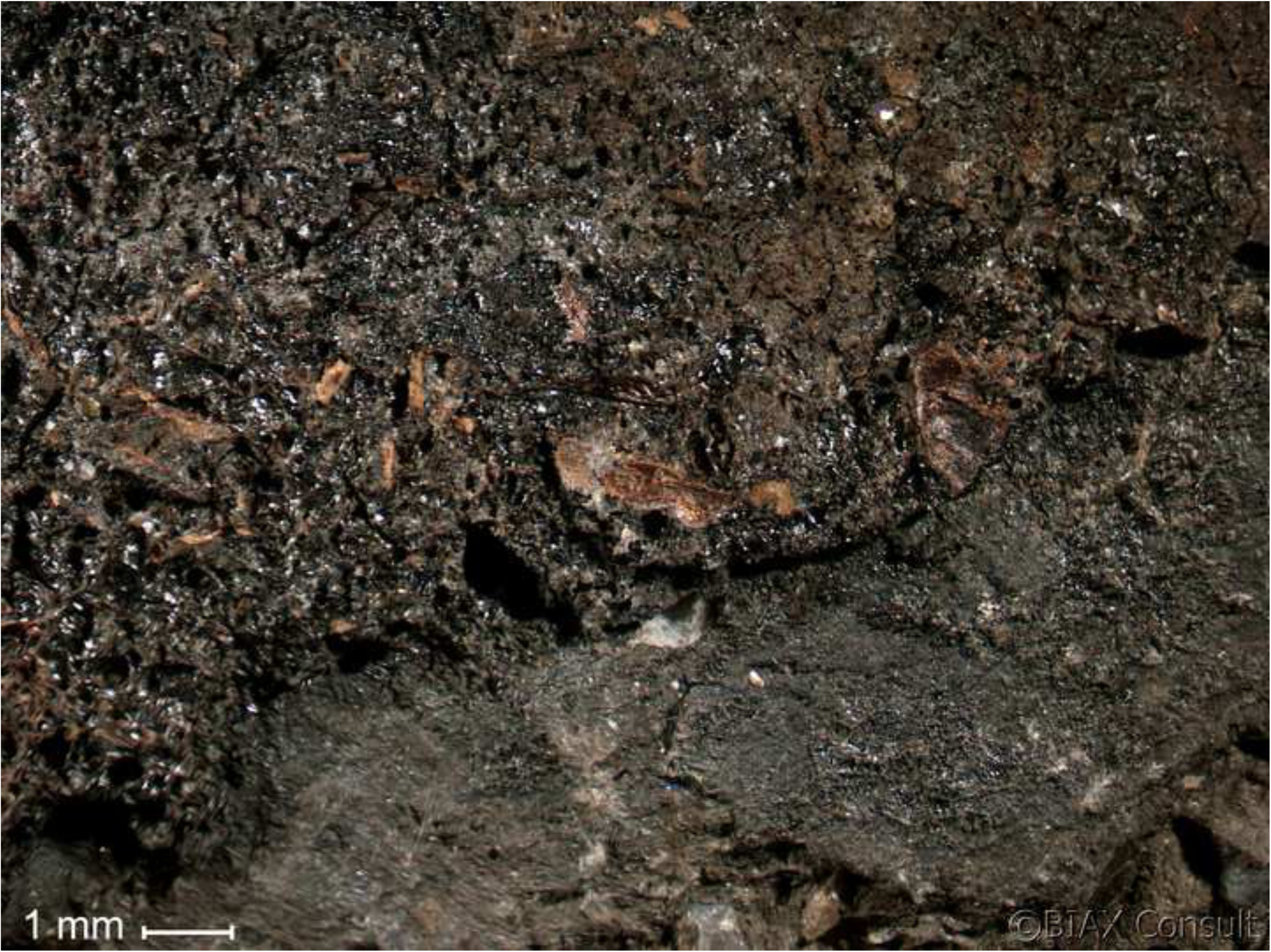
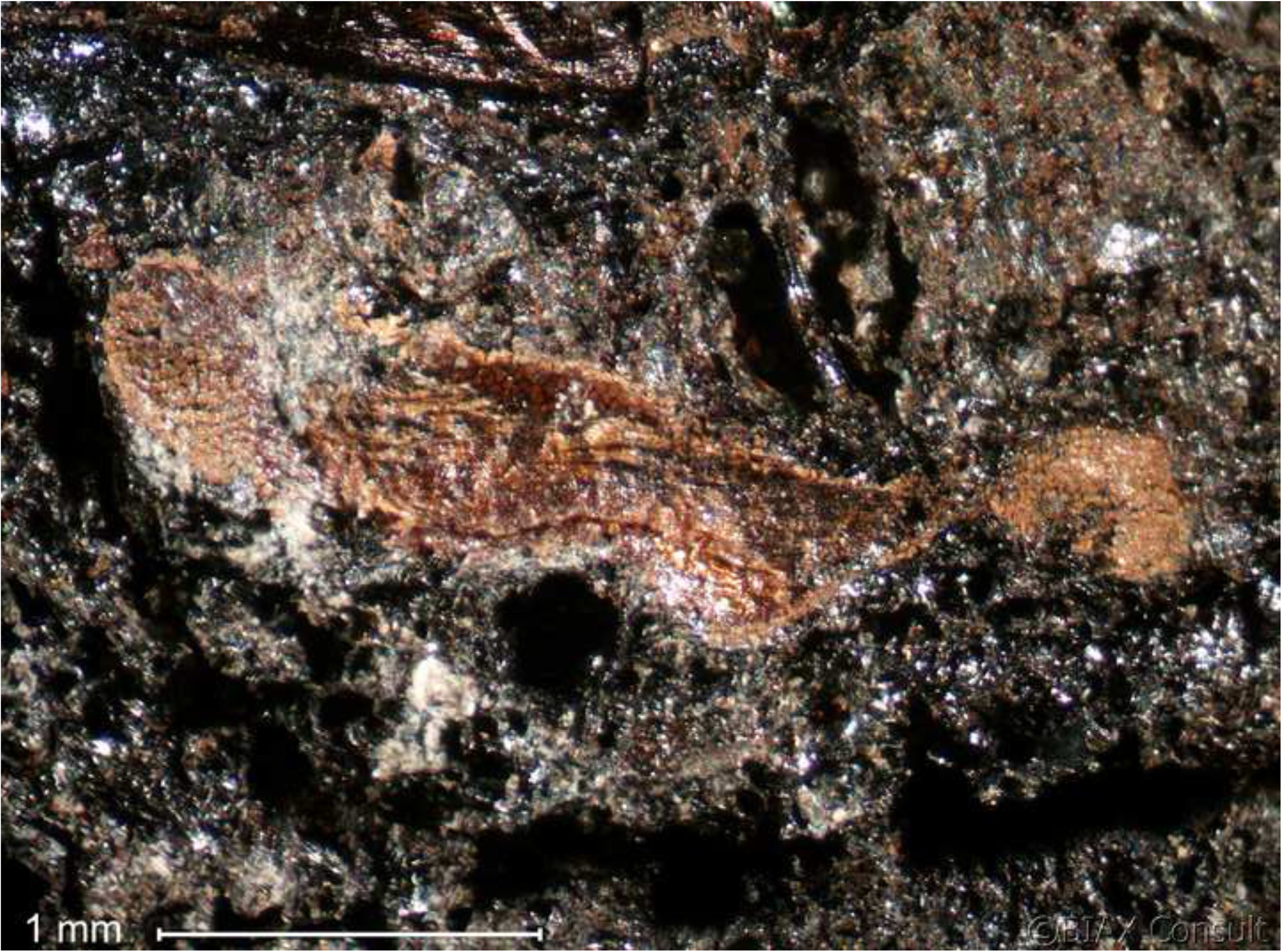
Oldenburg LA 77, sherd OLD 06, binocular microscope images of the upper face of the surface residue, showing overview (left) and detail of patches of cereal grain particles embedded in the residue matrix (right). Images: BIAX/M. van Waijjen.

**Fig 8.**
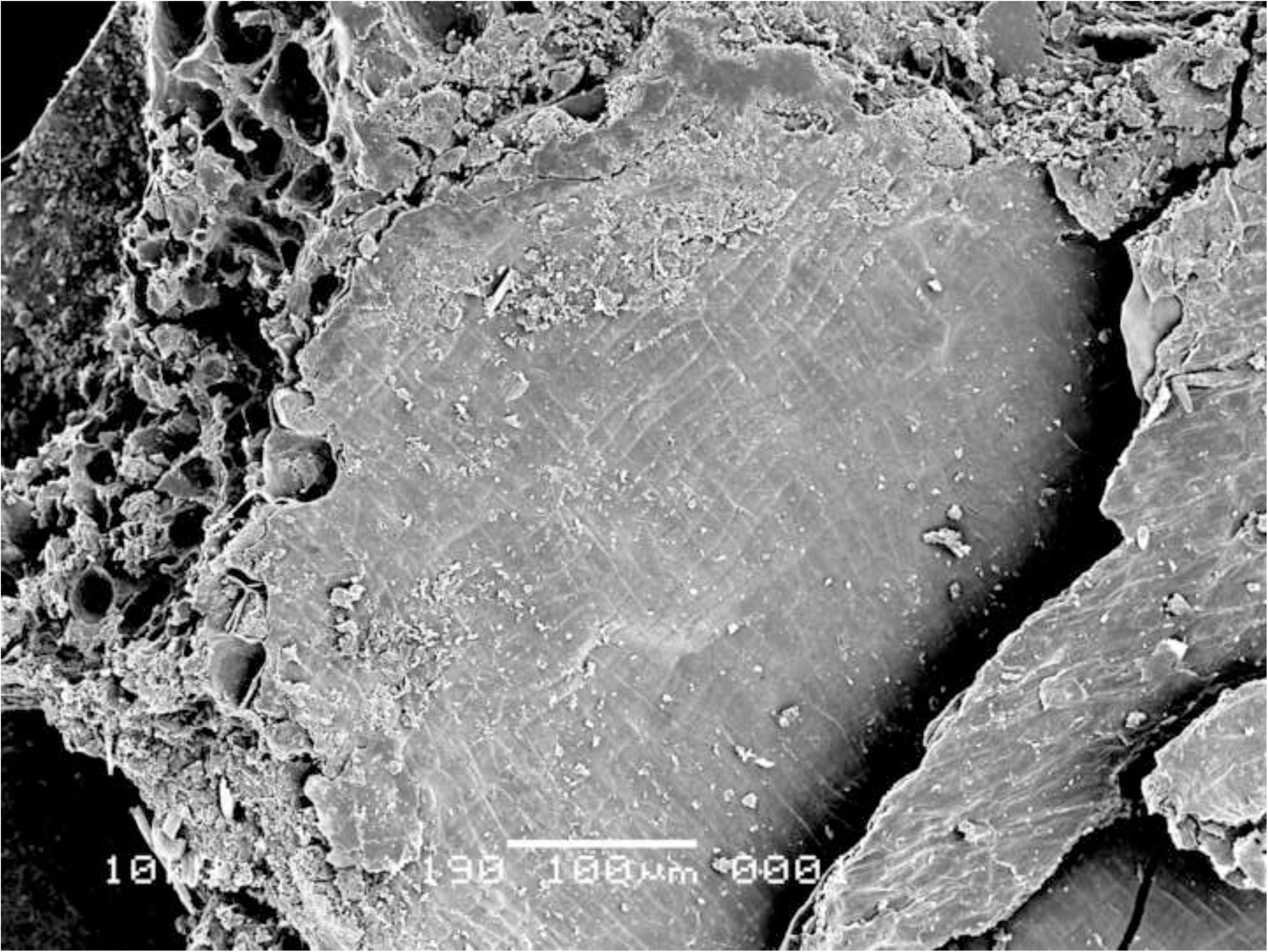
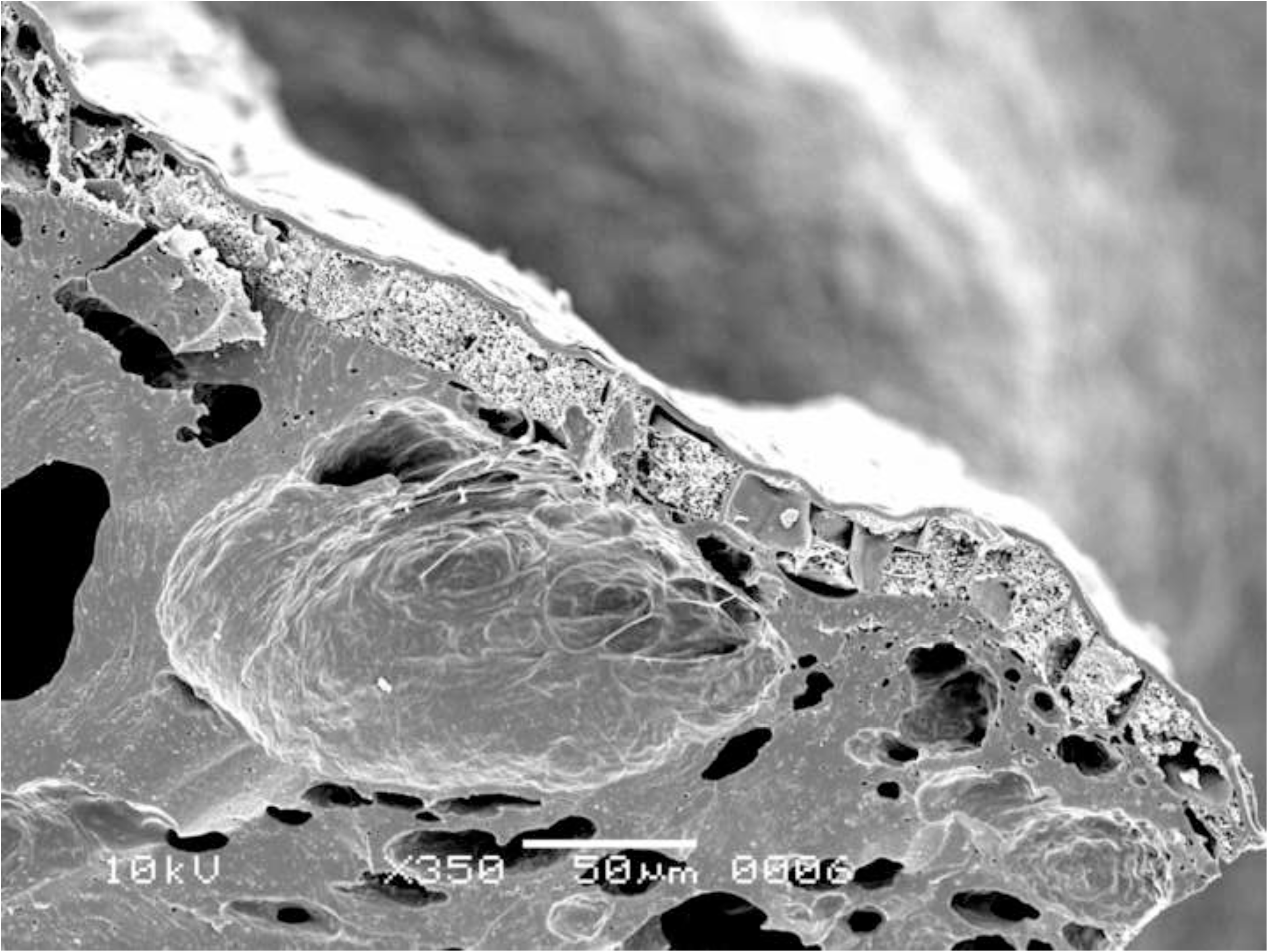
Oldenburg LA 77, sherd OLD 06, SEM images of grain fragments embedded in residue matrix, showing pericarp tissue (left) and single-layered aleurone accompanied by solid/fused (due to charring) grain starchy endosperm (in transverse view, right) of *Triticum* (here most likely emmer). Images: L. Kubiak-Martens.

**Fig 9.**
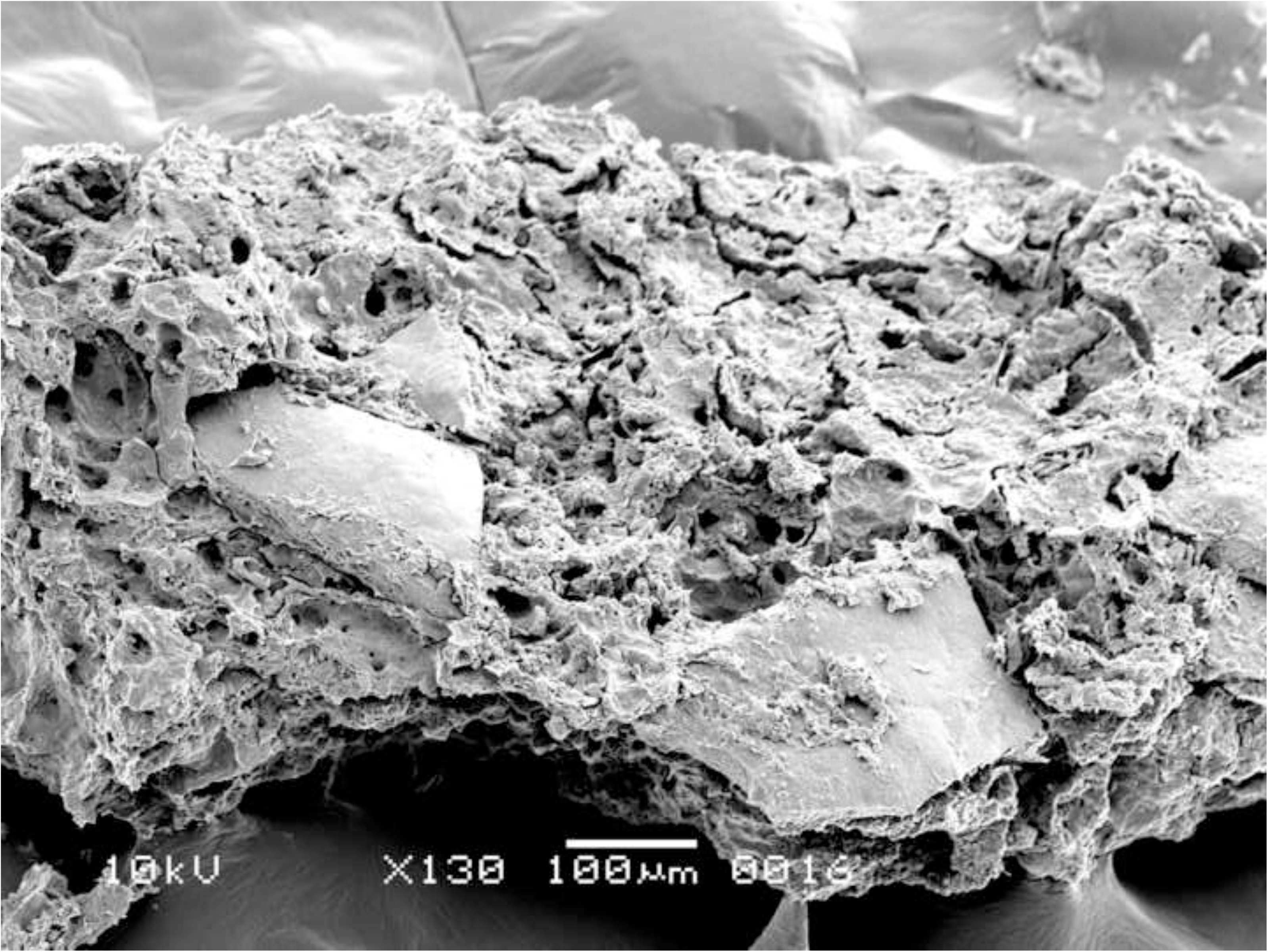
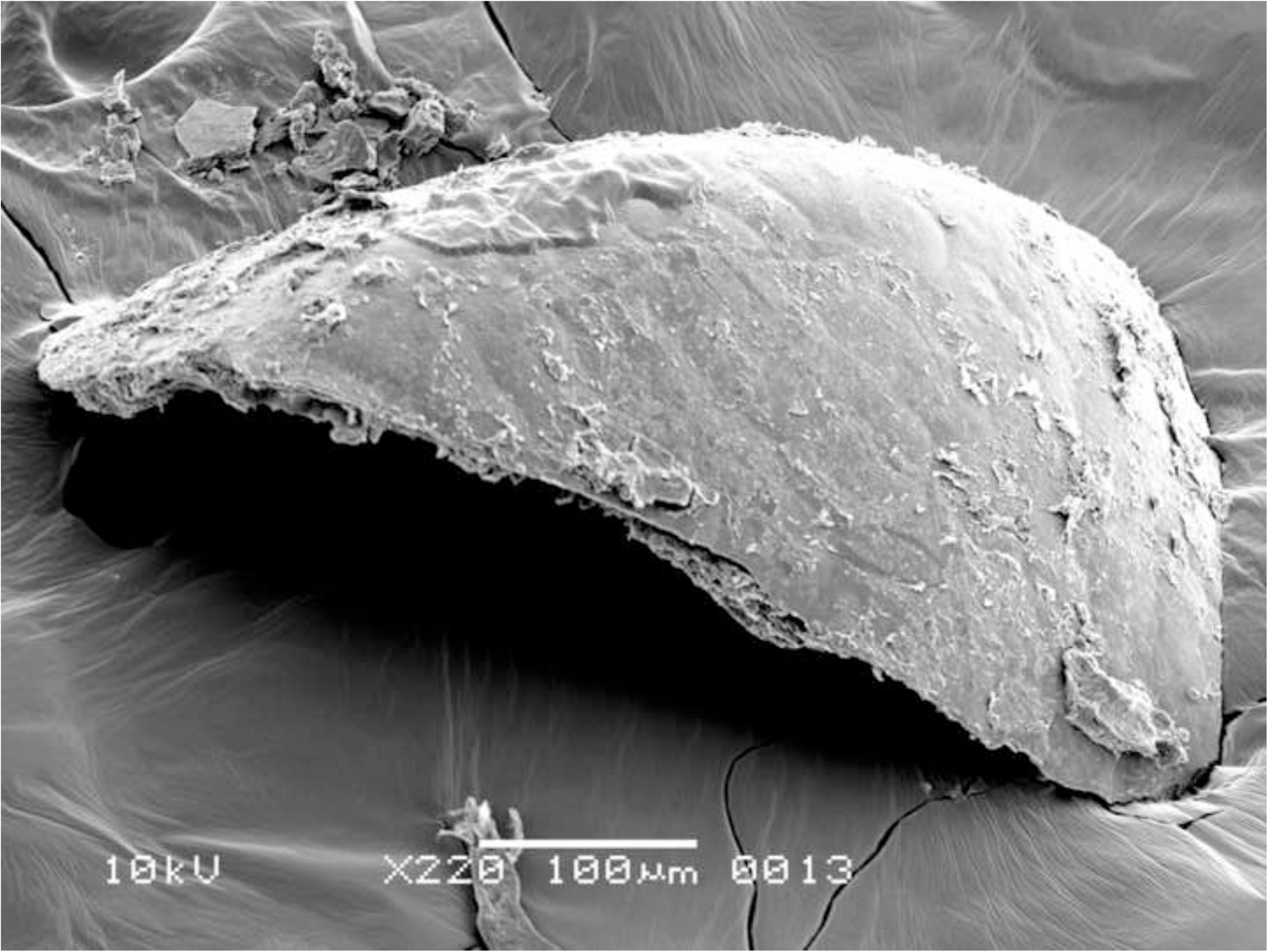
Oldenburg LA 77, sherd OLD 06, SEM images showing *Chenopodium album* seed fragments embedded in residue matrix (left) and *Chenopodium album* seed fragment isolated from the matrix for SEM photography (right). Images: L. Kubiak-Martens.

The absence of emmer chaff in the residues is noteworthy. In previous studies of food crusts from various Neolithic sites in the Netherlands, emmer light chaff particles (most likely of the lemma or palea) were frequently discovered in residues, suggesting that emmer must have been used in cooking. The presence of light chaff in the food crusts was attributed to the fact that despite dehusking of emmer, some light chaff would inevitably find its way into cooking pots [21, 32-33].

The aleurone cells of the emmer grain particles in Oldenburg residues have conspicuously thin walls. To determine the thickness of these cell walls, the R-STUDIO software was used to manually measure the shortest distance connecting two adjacent aleurone cell lumina in the SEM images. The aleurone cell wall thickness in the residues was compared with that of 10 emmer grains recovered from the settlement layer, one of which showed morphological evidence for sprouting (Fig 10). The raw measurements are provided in S3 (S3 Table 1 and S3 Table 2); the mean and median measurements are given in Table 2 and Fig 11. The mean and median values for the aleurone cell wall thickness in emmer grains retrieved from the Oldenburg food crusts are on average 1 μm (usually between 0.4 and 1.4 μm; in a few cases, a maximum thickness of about 1.6 μm was noted). Similar measurements were obtained for emmer grains from the occupation layer, which also display aleurone cell wall thinning. They show a mean and median thickness of approximately 1 μm and a maximum thickness of 1.5 μm. In contrast, the aleurone cell walls from archaeological emmer grains obtained from prehistoric sites, which are here used as a reference, show a thickness of c. 2 μm and, as such, do not display any markers for aleurone cell wall thinning [21]. The measurements obtained for aleurone cell wall thickness from both the crusts and the grains from Oldenburg correspond well with those reported for what are argued to be sprouted barley and emmer grains in earlier archaeobotanical studies, in particular, those conducted by Heiss and colleagues in which they suggested a threshold value for sprouted grains within the range of a mean/median of 1 µm for double cell wall thickness, and maximum value not exceeding 1.5 μm [34].

**Fig 10.**
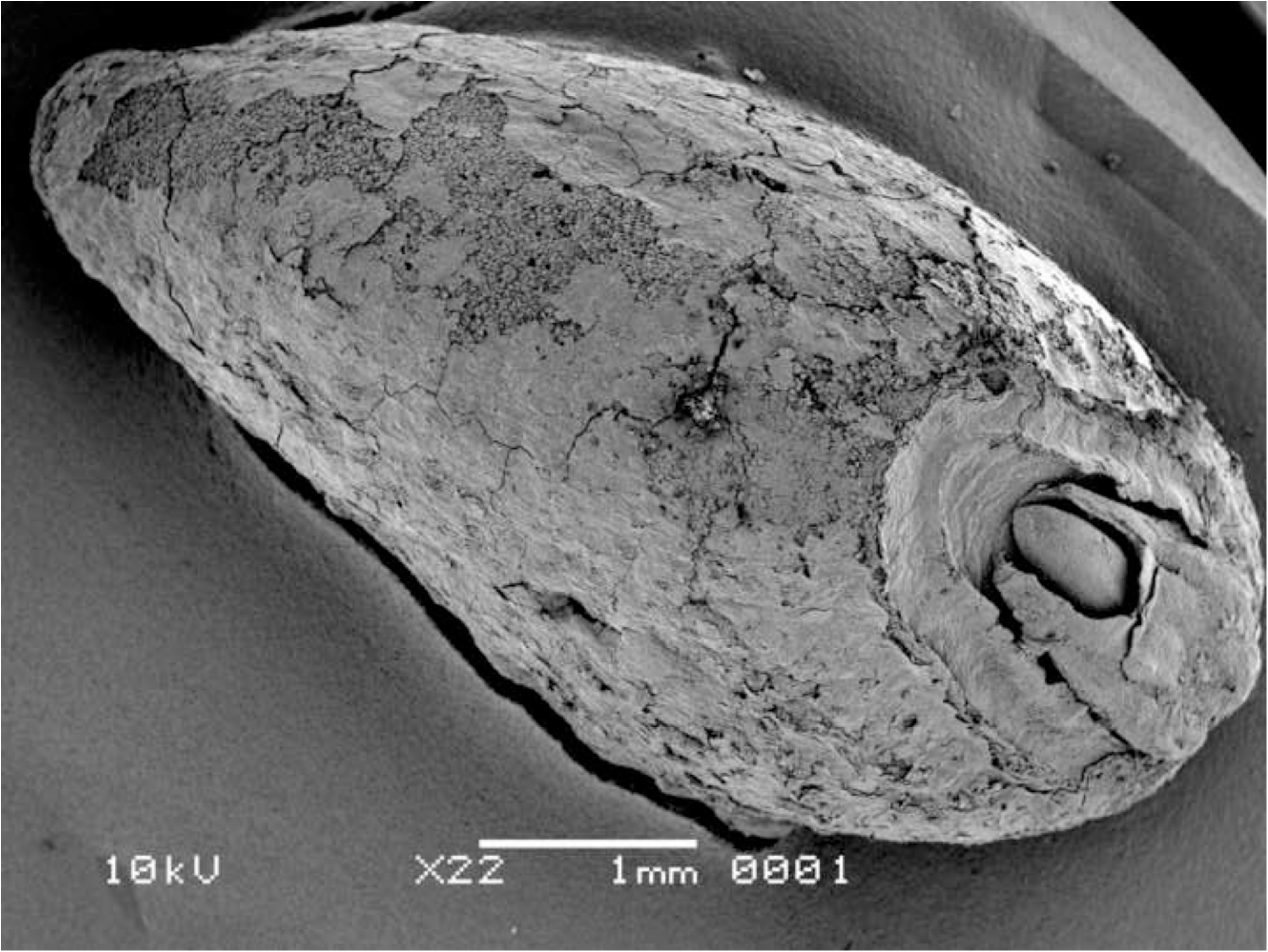
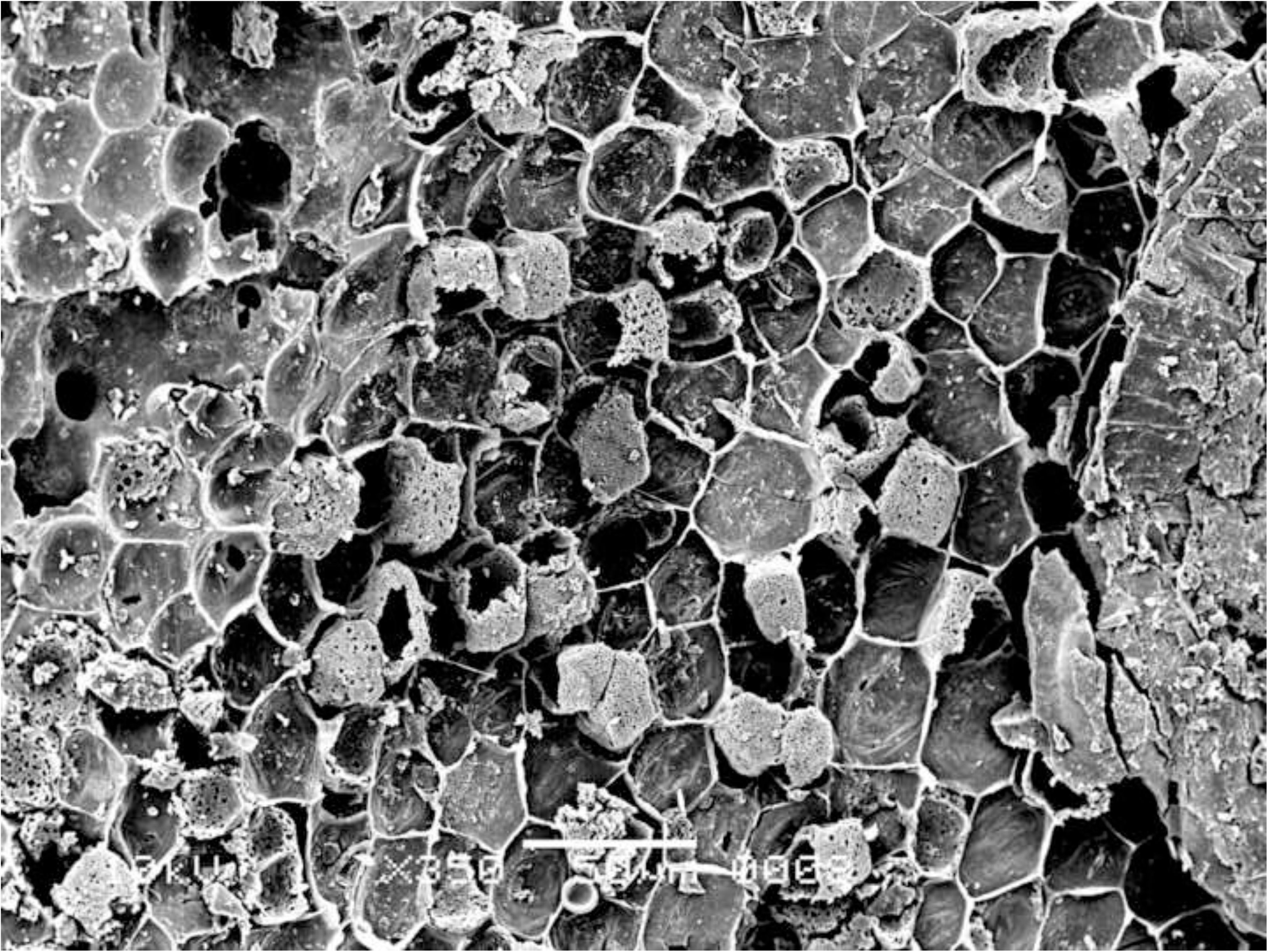
Oldenburg LA 77, charred emmer grain from the occupation layer, showing sprouted embryo (LA77-SED28-sp1) (left) and single-layered aleurone (LA77-SED27-sp6) with thinned aleurone cell walls in grain with no morphological evidence for sprouting (right). Images: L. Kubiak-Martens.

**Fig 11.**
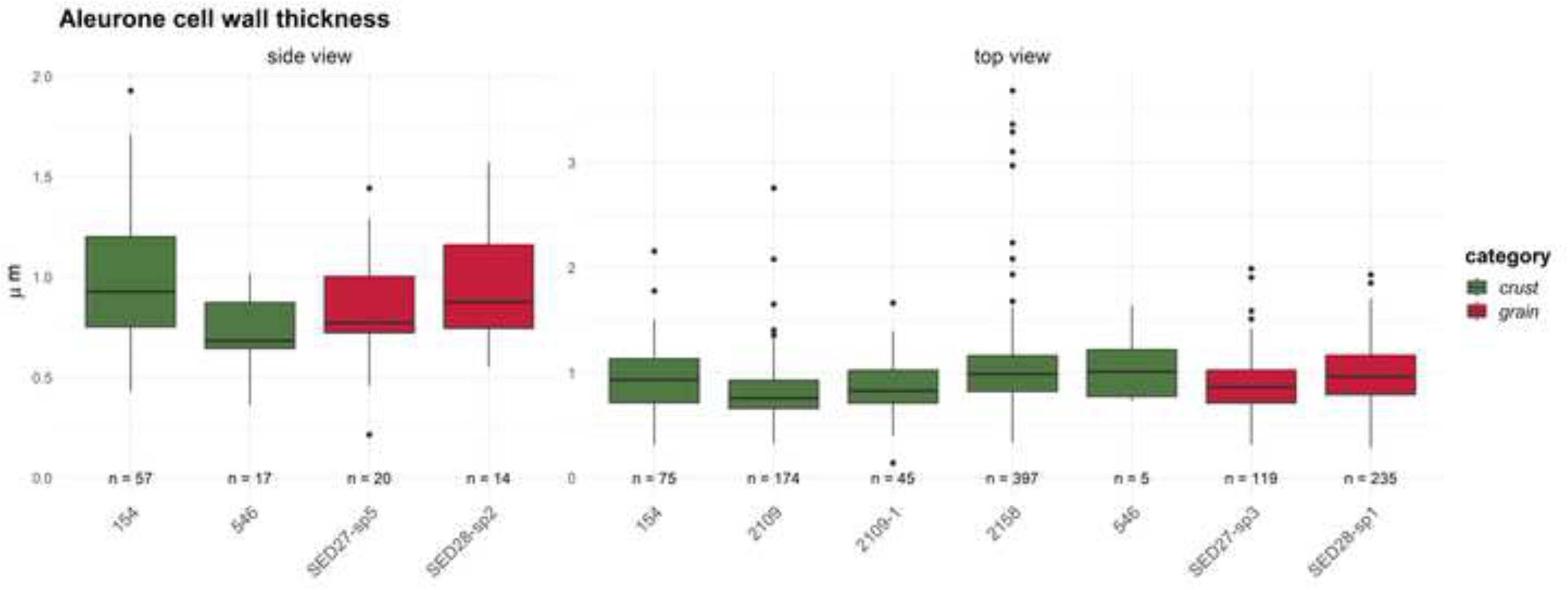
Oldenburg LA 77, aleurone double cell wall thicknesses in emmer grains (in μm) measured in food crusts, compared with the archaeological finds of charred emmer grains used in this study as reference material. Figure: BIAX/W. van der Meer.

**Table 2.**
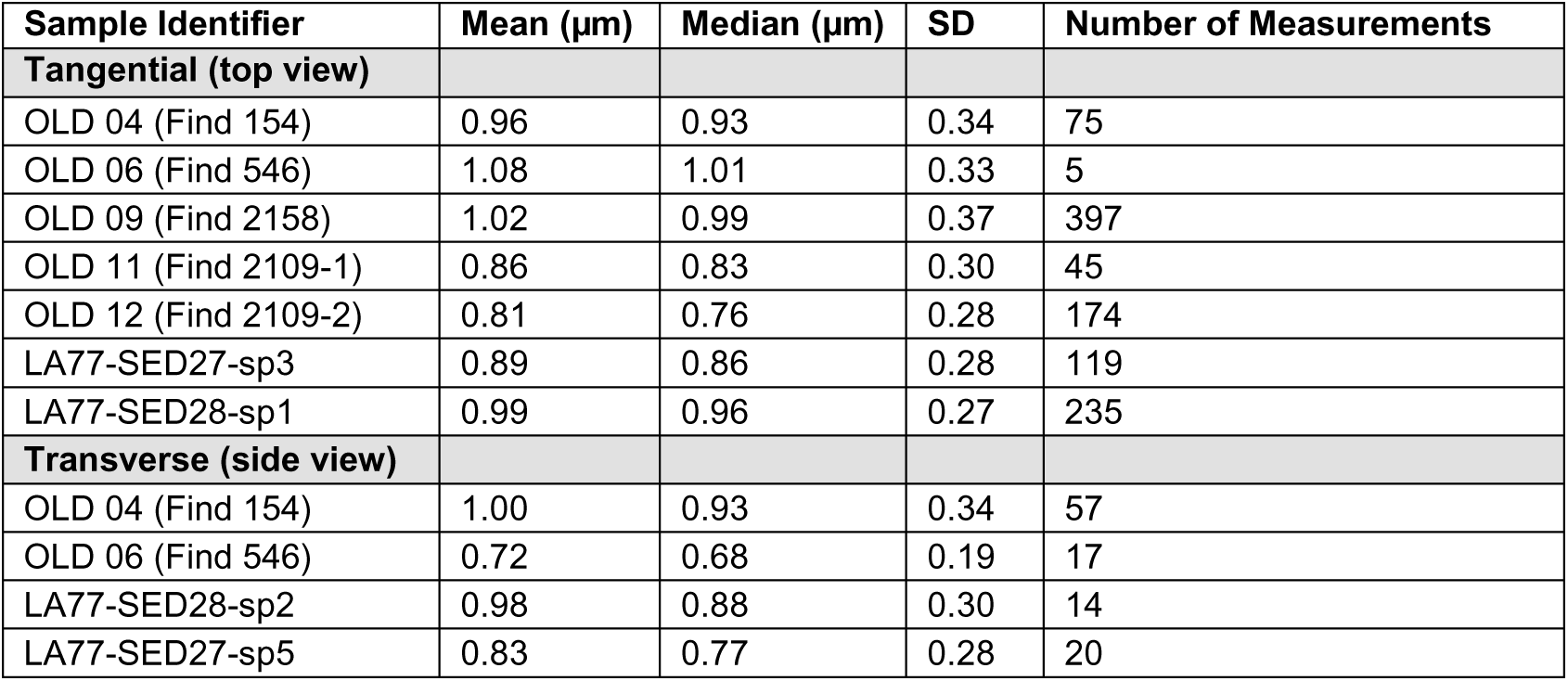
Oldenburg LA 77, mean and median values for aleurone double cell wall thickness in food crusts (identifiers starting with ‘OLD’) and two emmer grains from the settlement layer (identifiers starting with ‘LA77’). Data provided by BIAX/W. van der Meer. SD = standard deviation.

| Sample Identifier | Mean ($\mu\text{m}$) | Median ($\mu\text{m}$) | SD | Number of Measurements |
| --- | --- | --- | --- | --- |
| **Tangential (top view)** |  |  |  |  |
| OLD 04 (Find 154) | 0.96 | 0.93 | 0.34 | 75 |
| OLD 06 (Find 546) | 1.08 | 1.01 | 0.33 | 5 |
| OLD 09 (Find 2158) | 1.02 | 0.99 | 0.37 | 397 |
| OLD 11 (Find 2109-1) | 0.86 | 0.83 | 0.30 | 45 |
| OLD 12 (Find 2109-2) | 0.81 | 0.76 | 0.28 | 174 |
| LA77-SED27-sp3 | 0.89 | 0.86 | 0.28 | 119 |
| LA77-SED28-sp1 | 0.99 | 0.96 | 0.27 | 235 |
| **Transverse (side view)** |  |  |  |  |
| OLD 04 (Find 154) | 1.00 | 0.93 | 0.34 | 57 |
| OLD 06 (Find 546) | 0.72 | 0.68 | 0.19 | 17 |
| LA77-SED28-sp2 | 0.98 | 0.88 | 0.30 | 14 |
| LA77-SED27-sp5 | 0.83 | 0.77 | 0.28 | 20 |

Several additional factors may affect the thickness of aleurone cell walls in archaeological finds of grains, including infestation by mycelia fungi, endogenous enzyme activity, and thermal degradation [34]. However, in the case of the Oldenburg specimens, there was no evidence of a fungal infestation, such as the presence of hyphae or perforated cell walls.

Furthermore, many of the aleurone cells retained their cell content, which was intact, thus showing no evidence of endogenous enzyme activity. Extensive thermal degradation is also unlikely, as this would require cooking or charring temperatures exceeding 300°C [34], and in this study, the good preservation of polysaccharides points to heating or cooking at relatively low temperatures (see Chemical analyses section below).

### Food crust with milk-ripe barley

The SEM analysis of two of the surface residues, from OLD 09 (Fig 12) and OLD 10 (Fig 16), has revealed the presence of unripe or milk-ripe barley grains. The taxonomic identification process proved to be somewhat challenging due to the grains’ small dimensions, of approximately 4 mm x 1.4–1.8 mm (Fig 12 right and 13). However, the presence of multi-layered aleurone has been taken as the definitive diagnostic feature for the determination of their taxonomic identity (Figs 14, 15 and 17). In residue OLD 10, as many as three distinct cell layers of the aleurone tissue can be observed (Fig 17 right), clearly pointing to domesticated barley (*Hordeum vulgare*), which regularly develops a multi-layered aleurone, usually consisting of three layers of cuboidal cells [35].

**Fig 12.**
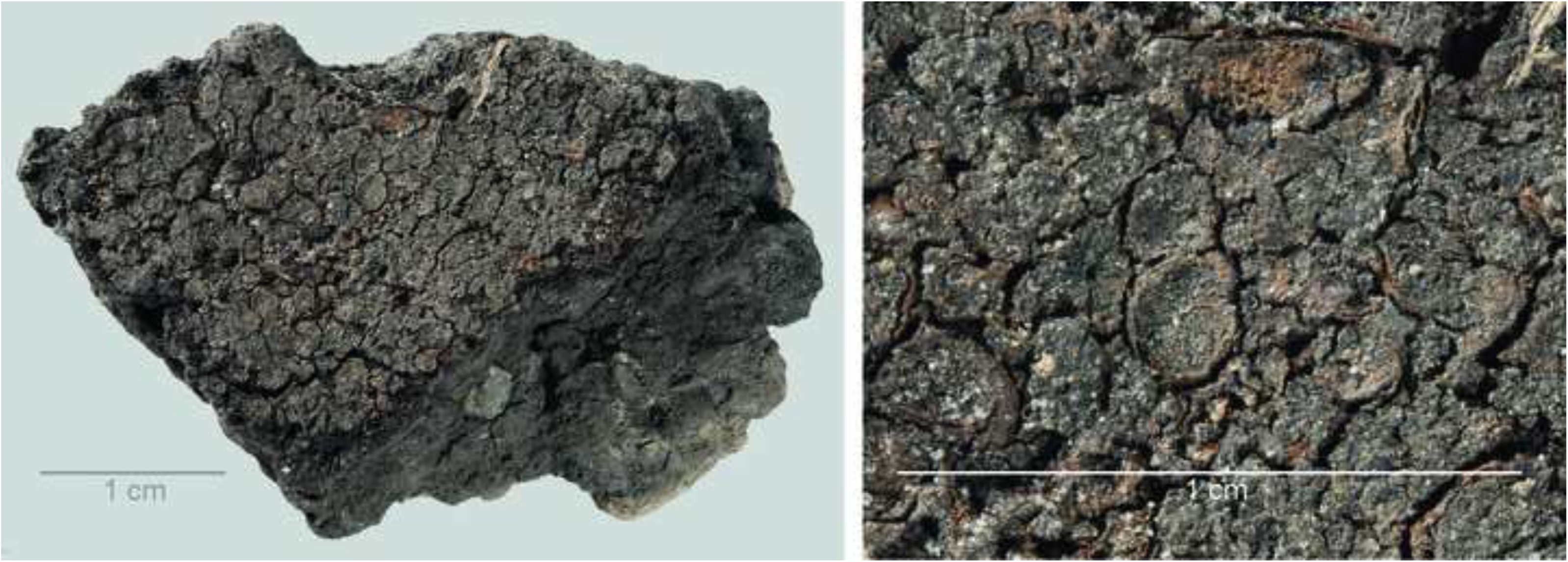
Oldenburg LA 77, sherd OLD 09, binocular microscope image of food crust in top view (left) and of a detail of the food crust with multiple immature barley grains (ca. 4 mm x 1.4–1.8 mm in size) embedded in the matrix (right). Images: Institut für Ur- und Frühgeschichte, Kiel.

**Fig 13.**
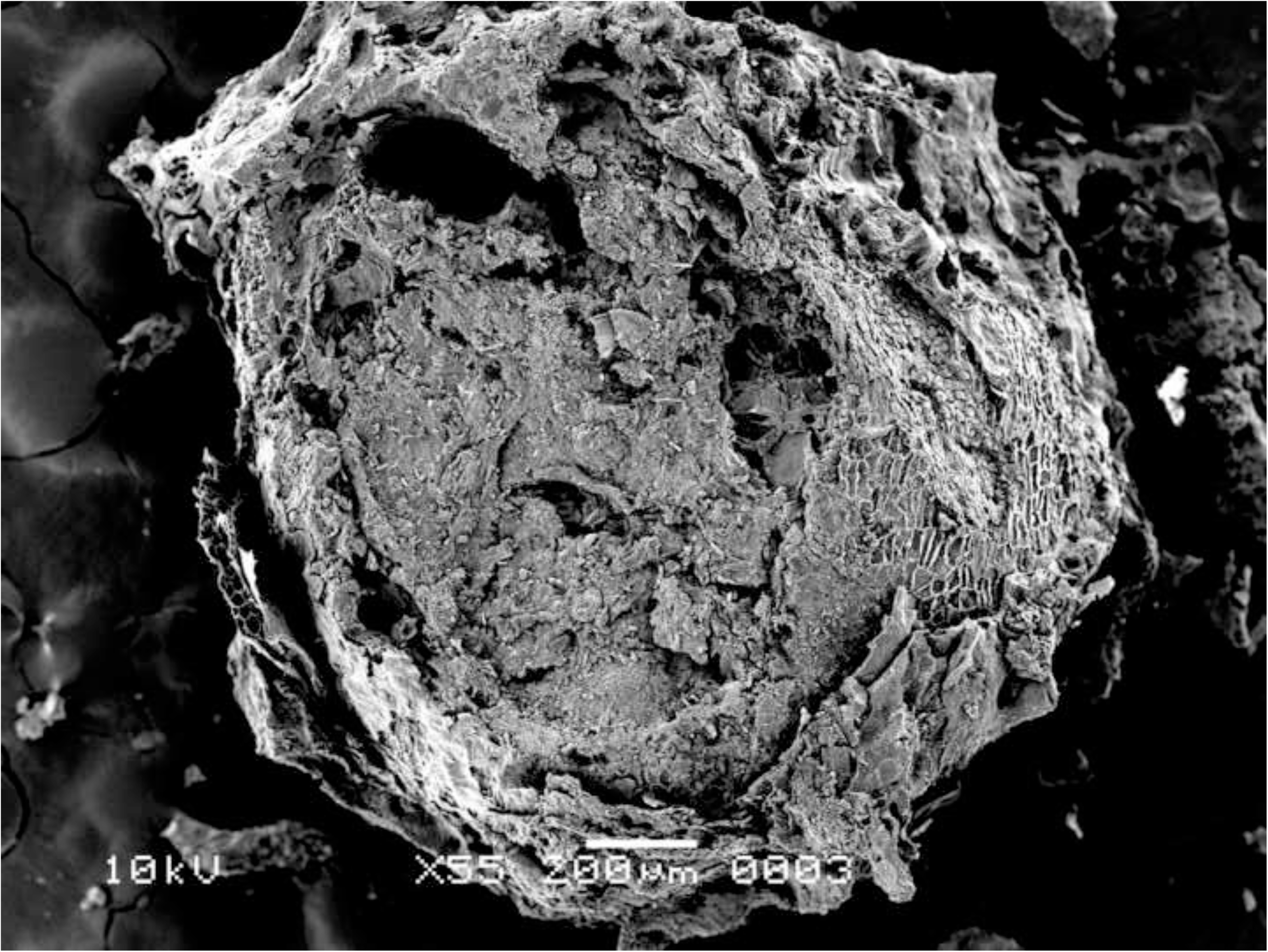
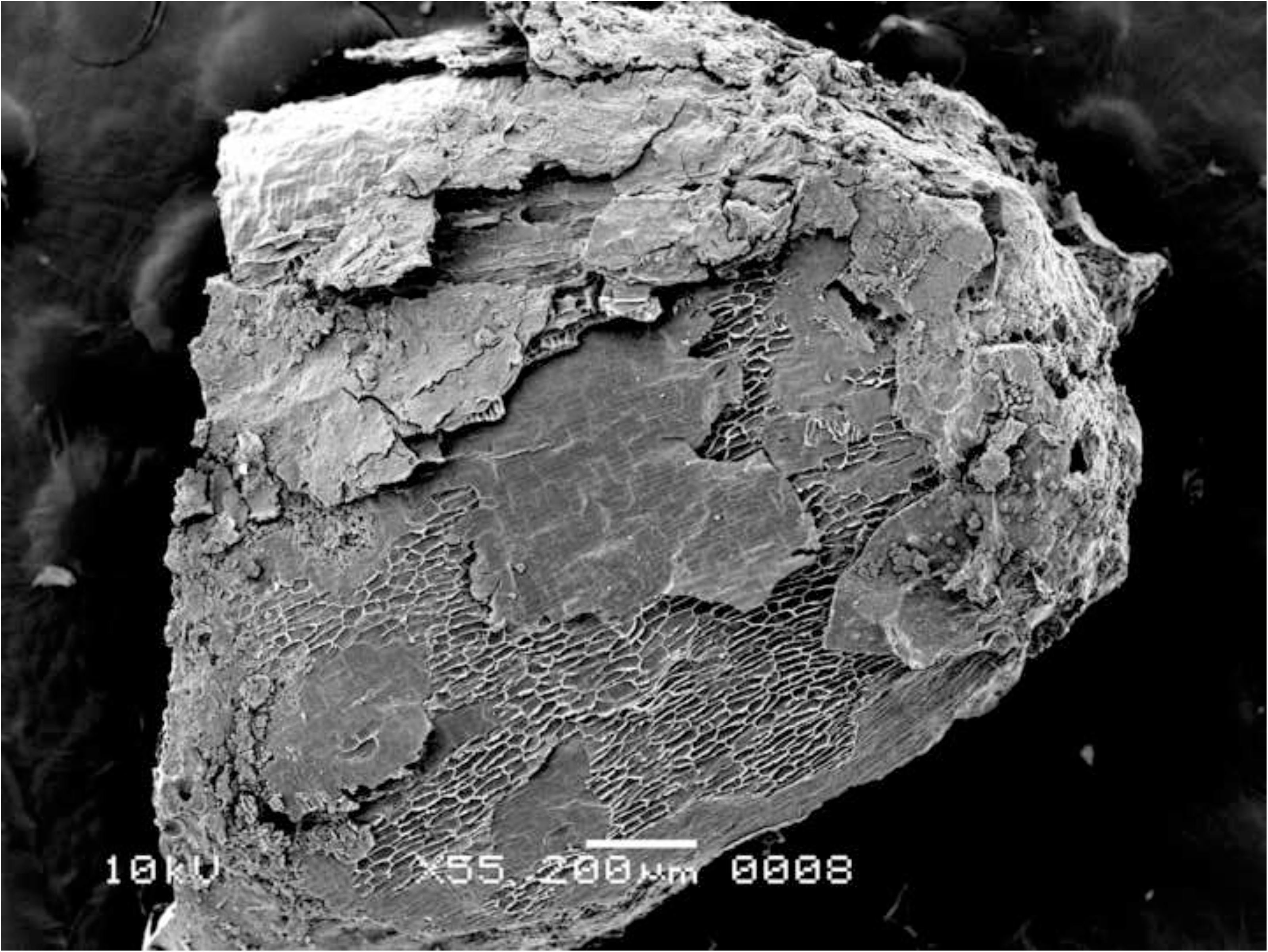
Oldenburg LA 77, sherd OLD 09, immature barley grains isolated from the crust for SEM photography, transverse view (left), and side view (right), showing partially preserved seed coat and pericarp. Images: L. Kubiak-Martens.

**Fig 14.**
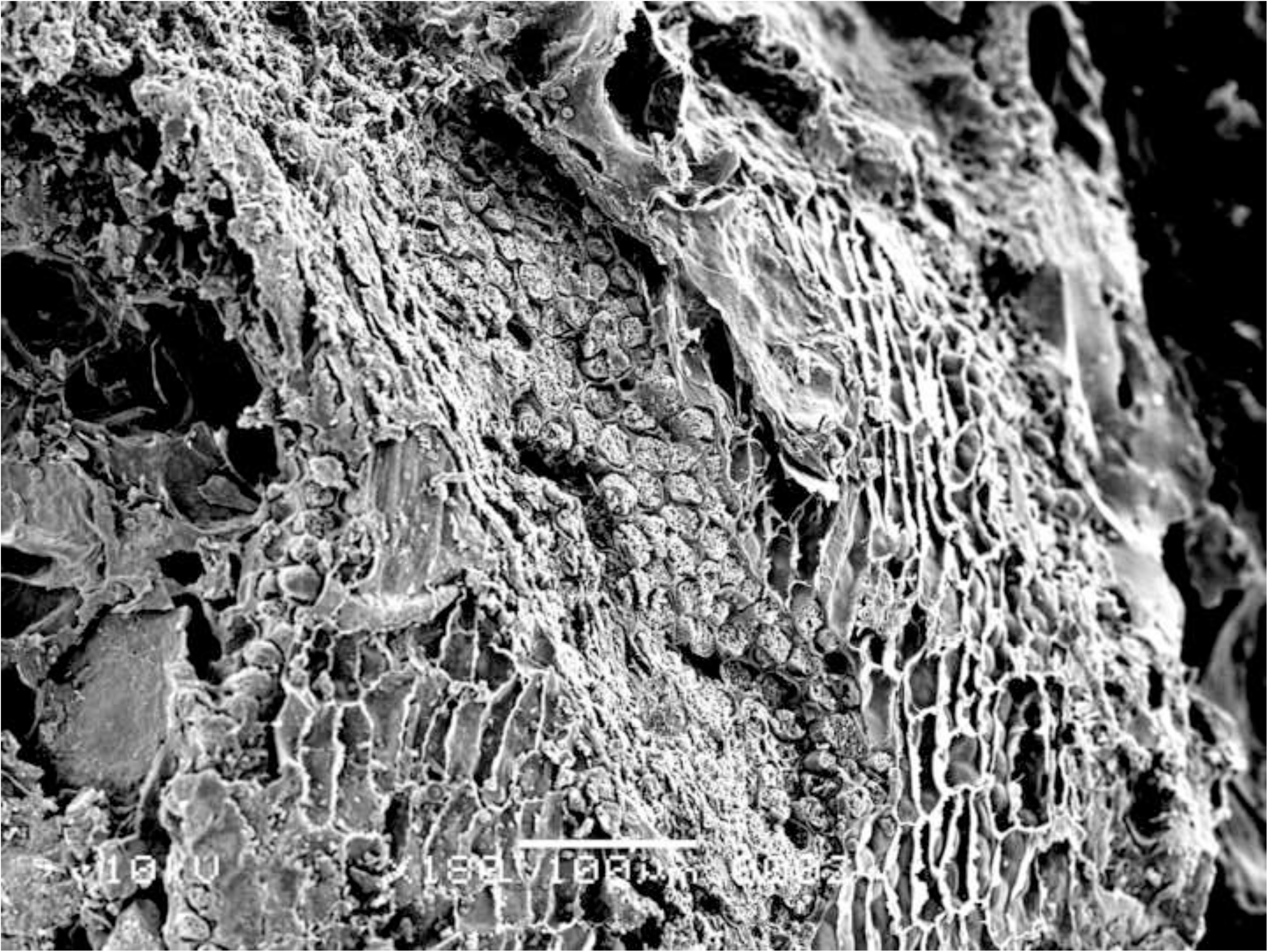
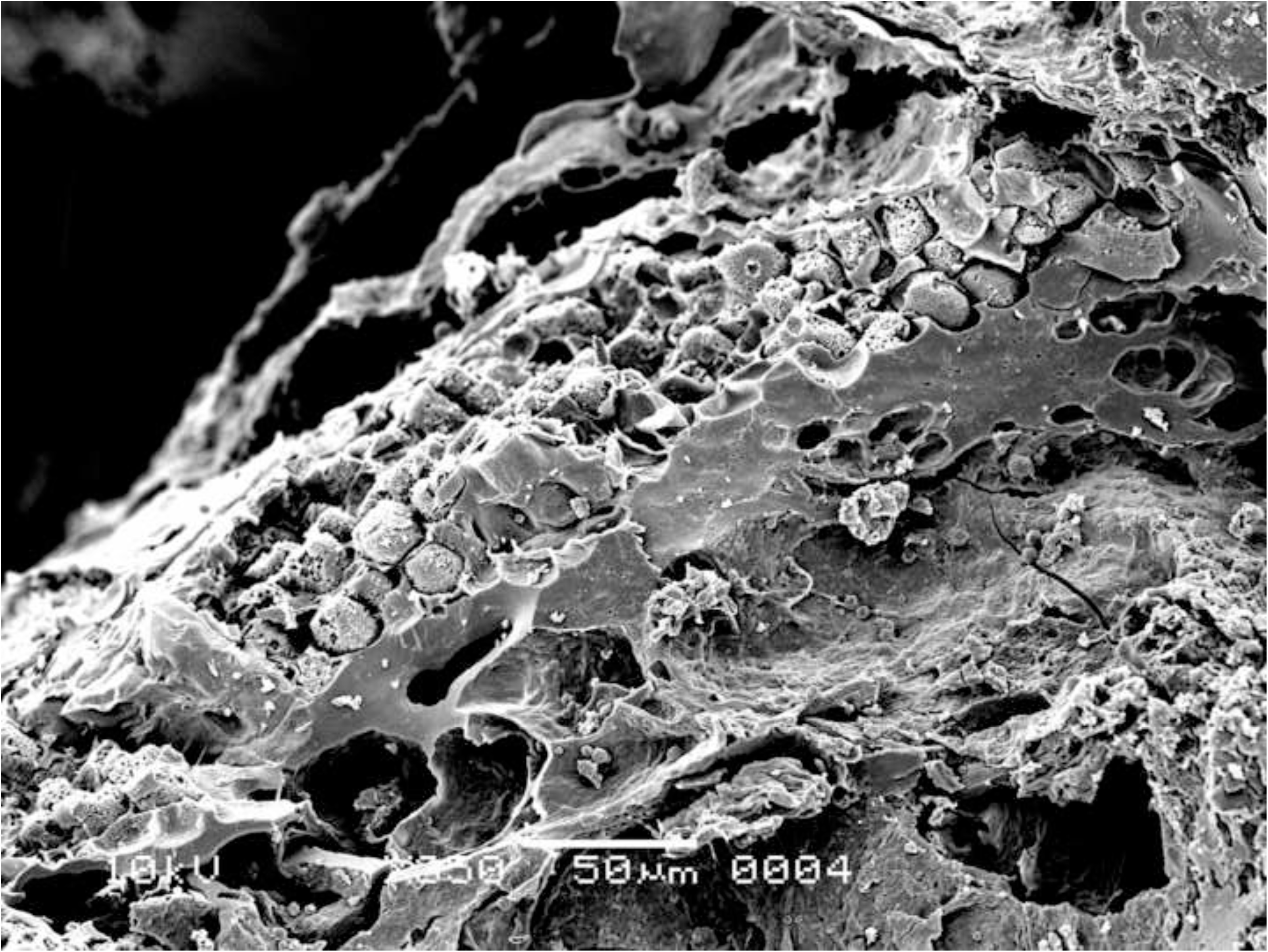
Oldenburg LA 77, sherd OLD 09, SEM images showing multicellular aleurone layer as observed in barley, accompanied by pericarp tissues, top view (left), and multicellular aleurone layer as observed in barley, transverse view (right). Images: L. Kubiak-Martens.

**Fig 15.**
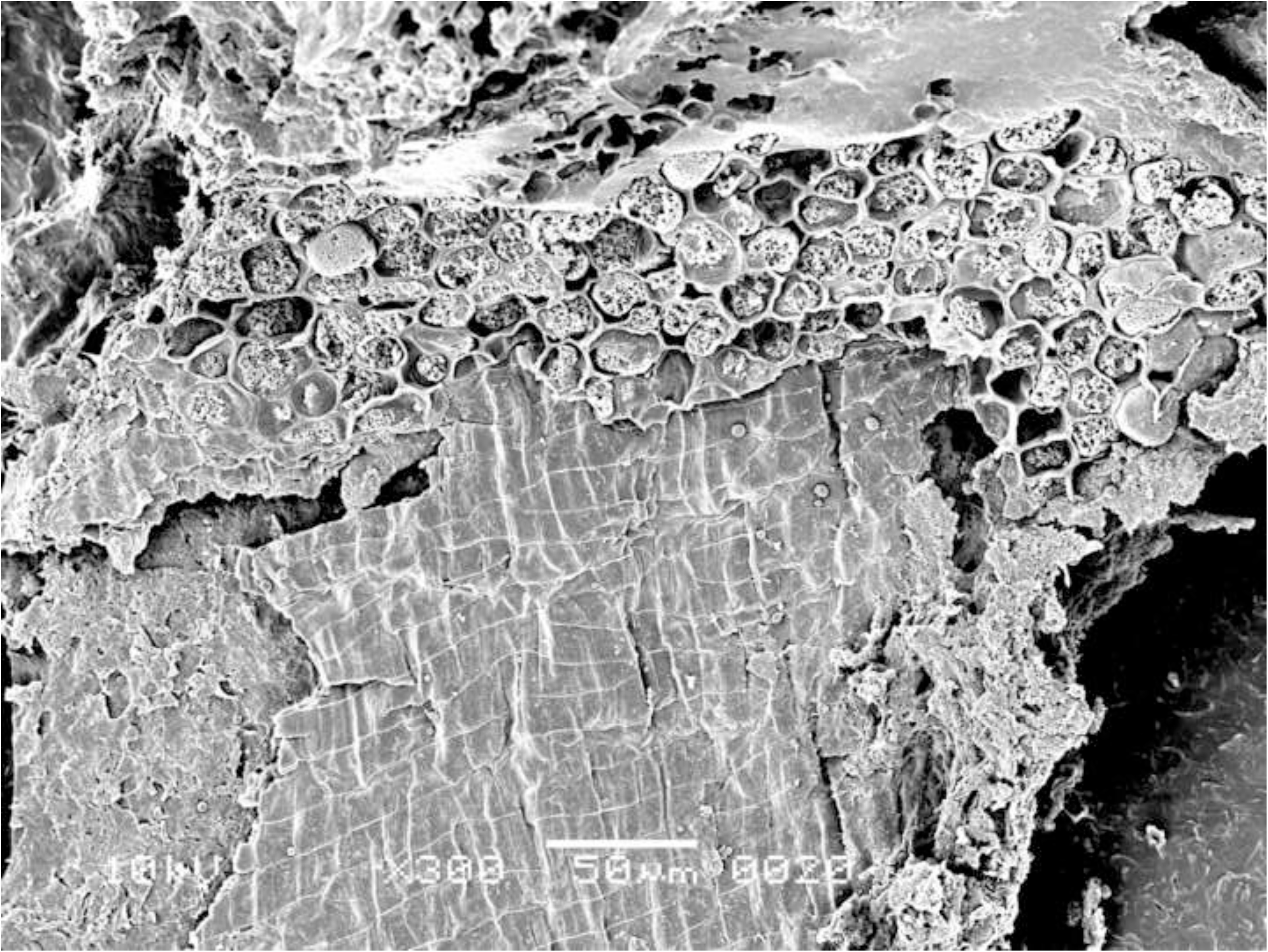
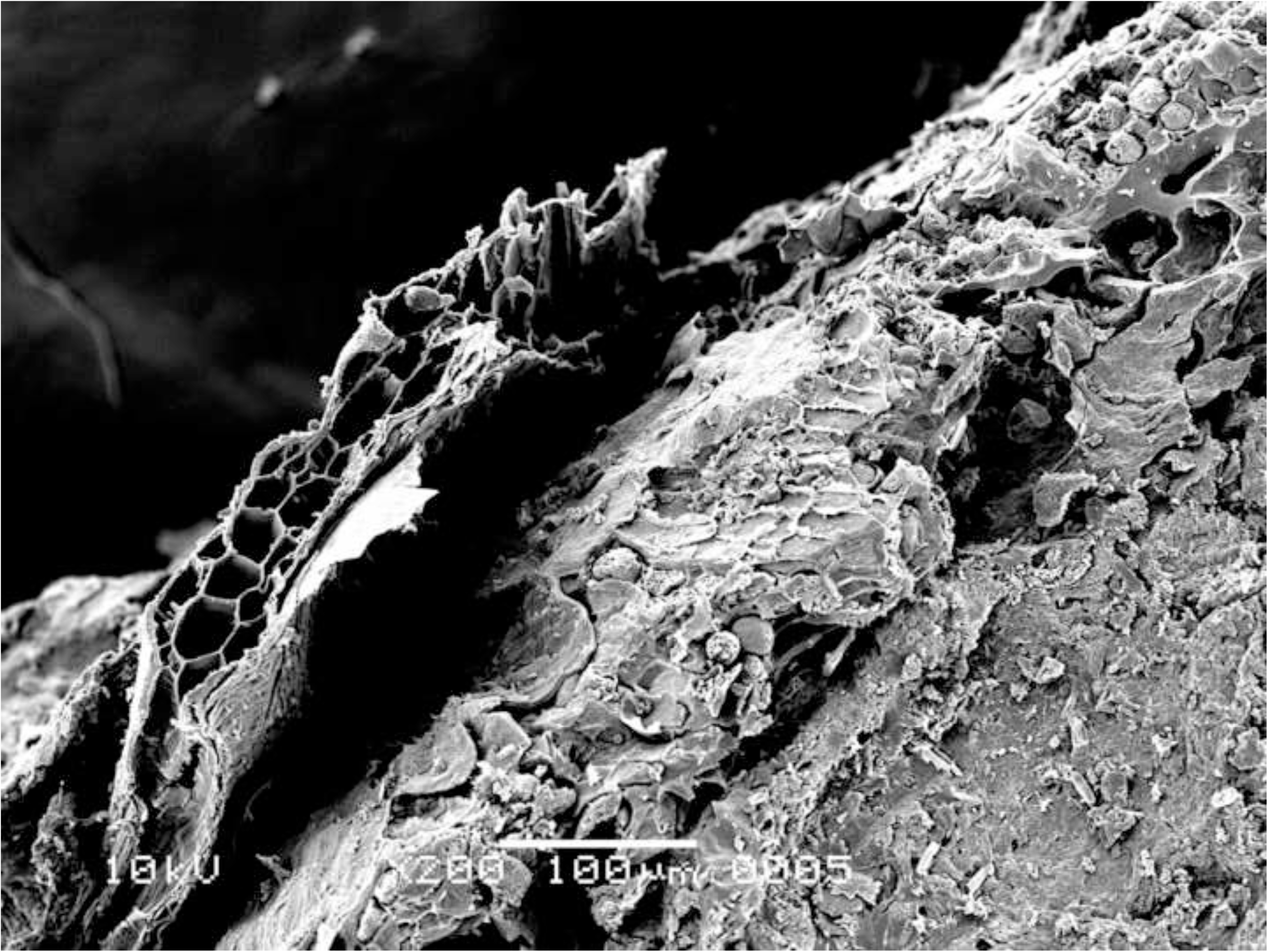
Oldenburg LA 77, sherd OLD 09, SEM images showing multicellular aleurone layer as observed in barley, accompanied by pericarp tissues, top view (left) and the edge of a barley grain fragment, revealing multicellular aleurone layer, pericarp tissues, and likely chaff attached to the pericarp (right). Images: L. Kubiak-Martens.

**Fig 16.**
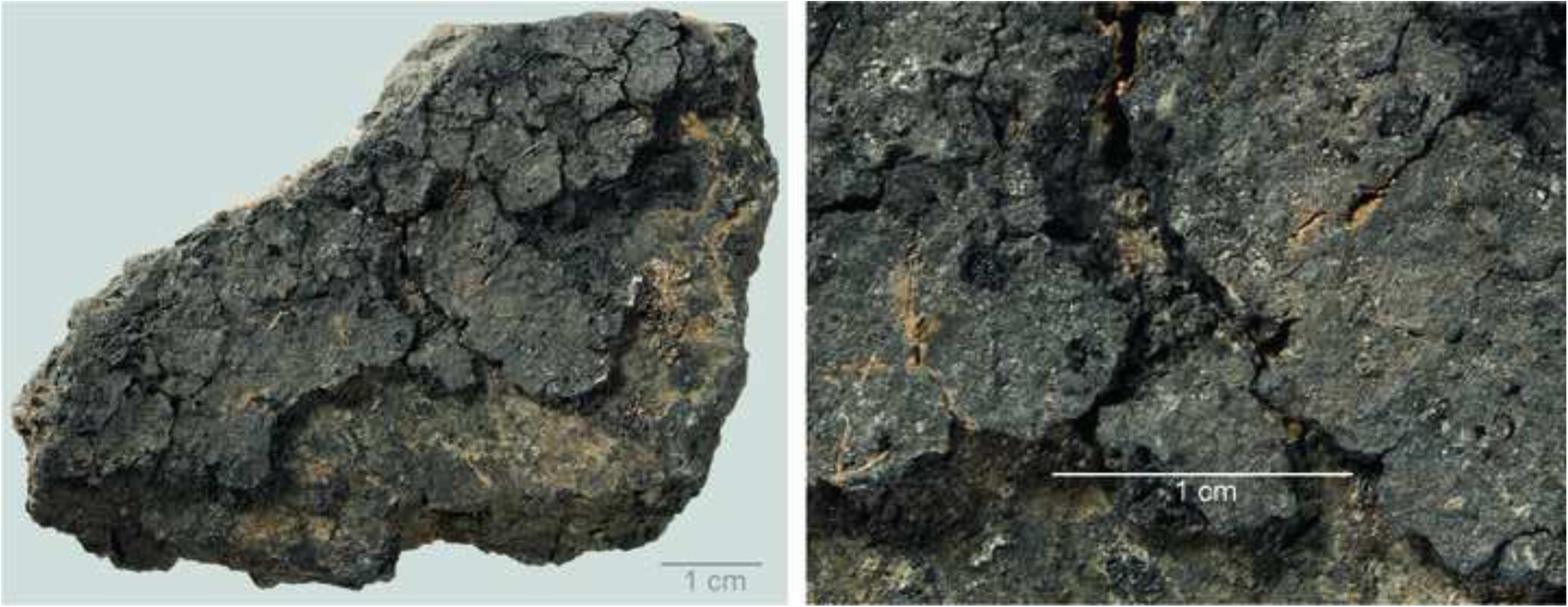
Oldenburg LA 77, sherd OLD 10 with food crust, overview (left), and detail showing immature barley grains embedded in the residue matrix (right). Images: Institut für Ur- und Frühgeschichte, Kiel.

**Fig 17.**
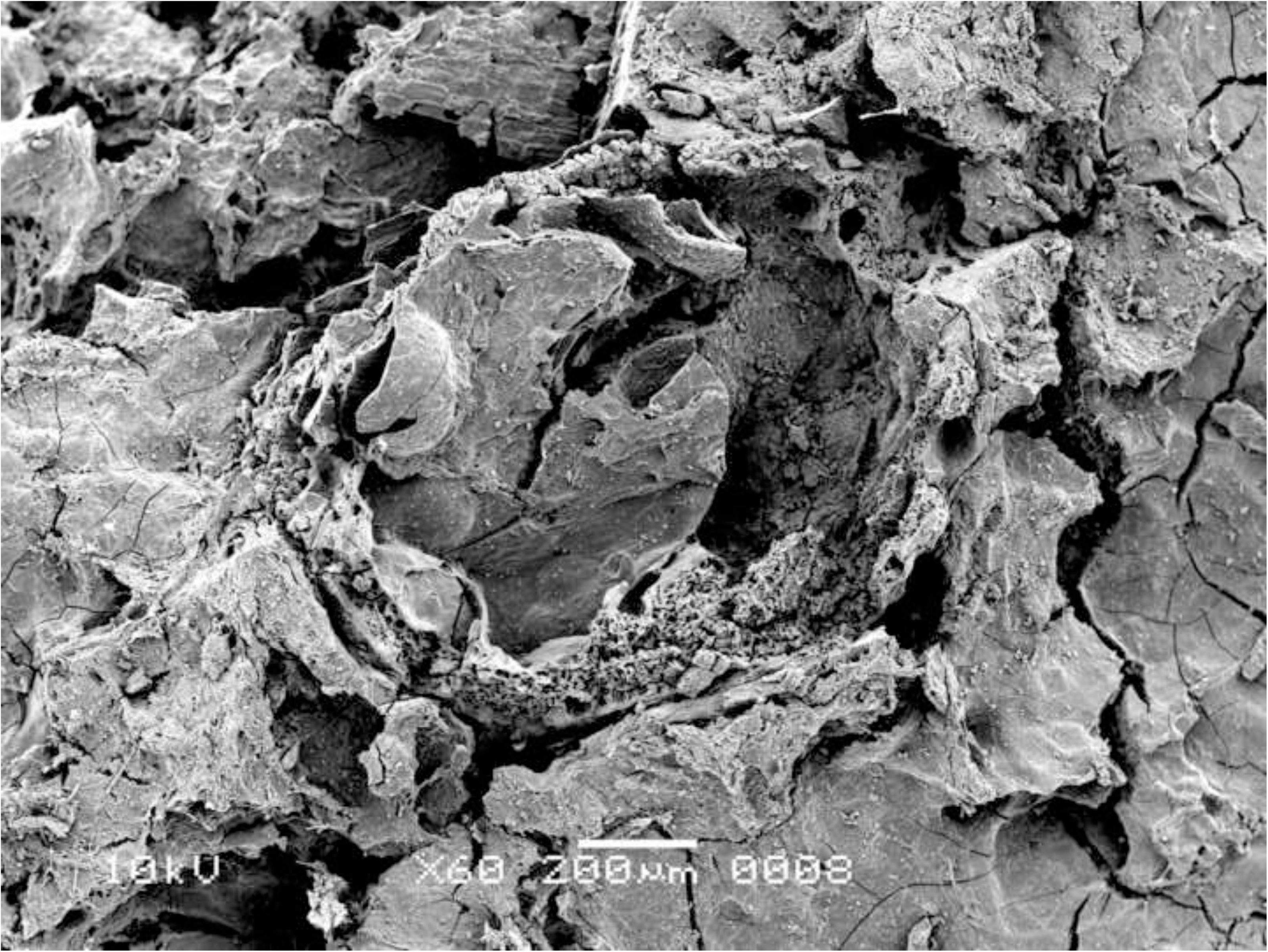
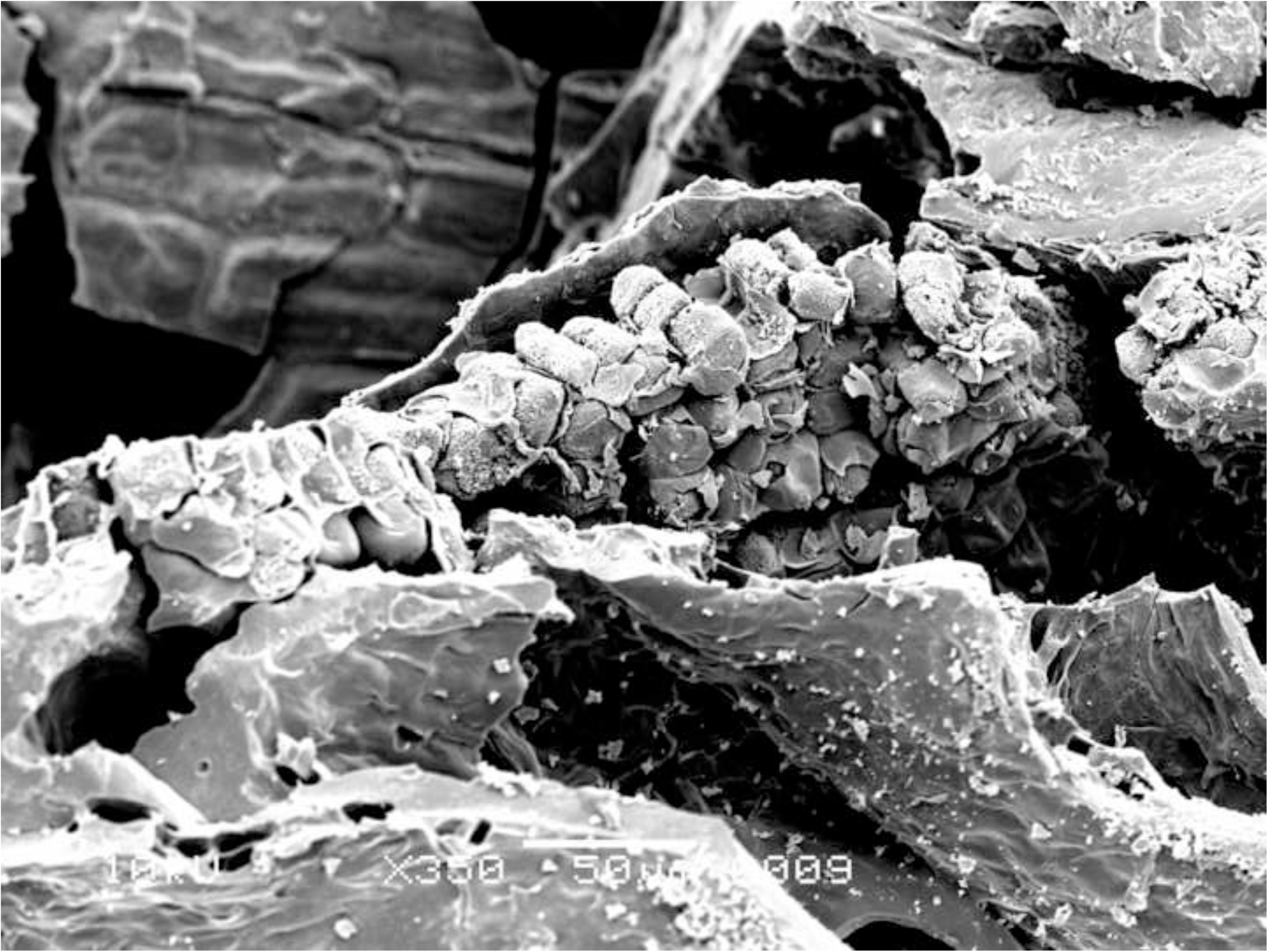
Oldenburg LA 77, sherd OLD 10, SEM images showing the outline of immature barley grain (left) and the multicellular aleurone characteristic of barley (right). The endosperm has collapsed during cooking or charring, due to the high moisture content of the immature grains. Images: L. Kubiak-Martens.

In addition to the small size of the barley grains, the presence of chaff, most likely lemma or palea, attached to the grains also suggests that they were harvested while the grain was not yet fully ripe (Fig 15 right). In the naked barley documented among the archaeobotanical remains from Oldenburg LA 77, the fully ripe grains had been freed from the chaff by threshing. But in the Oldenburg surface residues, light barley chaff was still attached to the grains, thus indicating the use of immature, green grains. Immature grains contain higher levels of moisture and lower levels of starch than mature grains. The endosperm portion of the unripe barley grains in OLD 10 has collapsed (Fig 17 left), likely during cooking or charring and likely due to the grains’ high moisture content. In contrast, the endosperm portion of the emmer grains discovered in the SEM analyses in this study is more solid and starchy, which is typical for cooked mature grains.

## Chemical analyses

### ATR**–**FTIR results

All residues (n=21) were measured in triplicate, and the samples showed fairly homogeneous chemistry. The results of the ATR–FTIR analyses are shown in detail in S4 and illustrated with FTIR spectra in Fig 18, while a summary can also be found in Table 3. Identification is based on earlier FTIR examination of heated and charred food remains [36-39] and relevant studies in archaeology and art conservation [40-41]. It is important to keep in mind that ATR-FTIR is a screening method and the analysis of complex organic materials only shows characteristics of components that form 5% or more of the volume of the mixture. Compounds that are present in very small proportions may not be detected.

**Fig 18.**
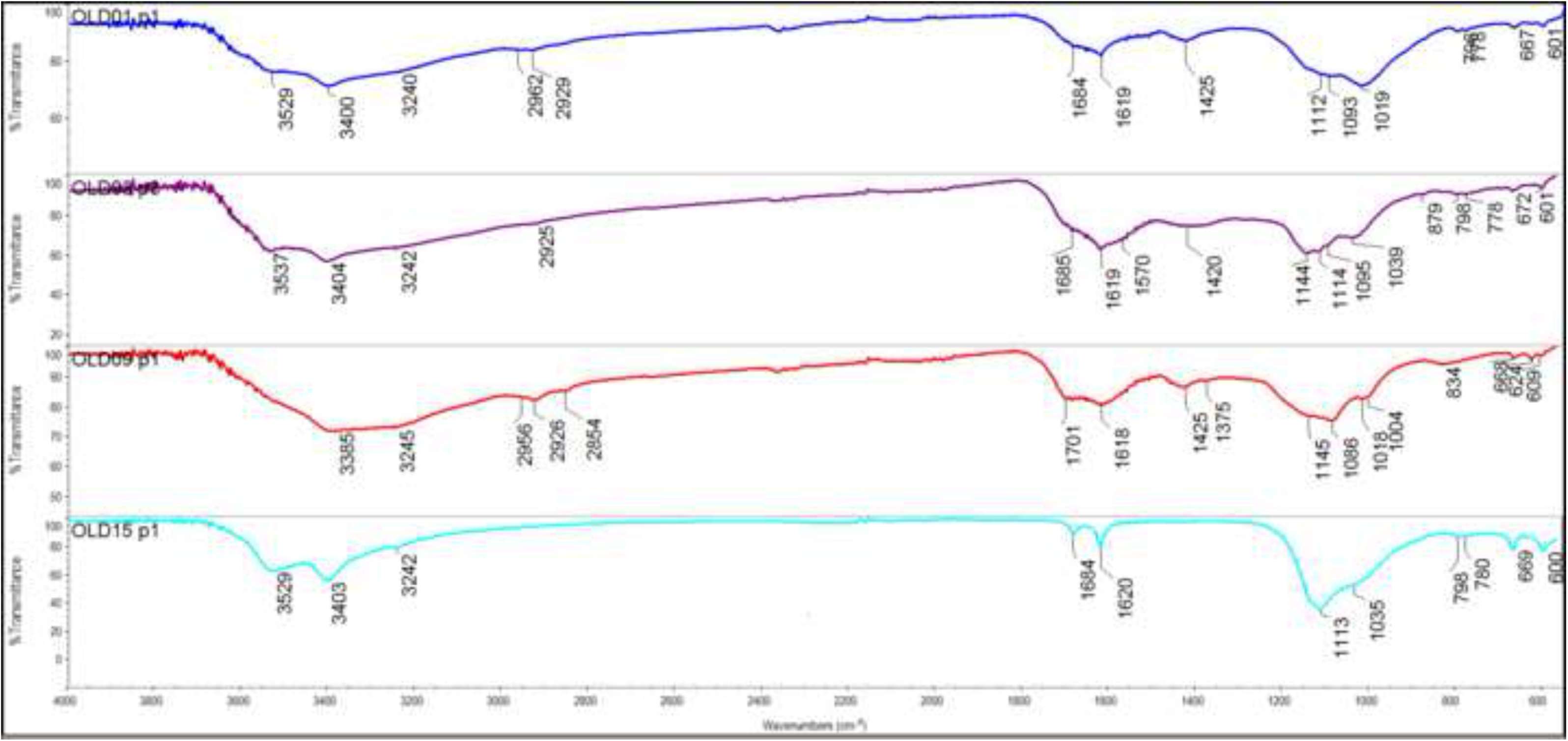
FTIR spectra, from top to bottom: OLD 01, 02, 09 and 15, showing the relative transmissions (in %) against the wave number (from 4000-550 cm^-1^). The first three spectra show characteristics typical of polysaccharides; the last spectrum is typical of calcium sulfate. Graph: T.F.M. Oudemans.

**Table 3.**
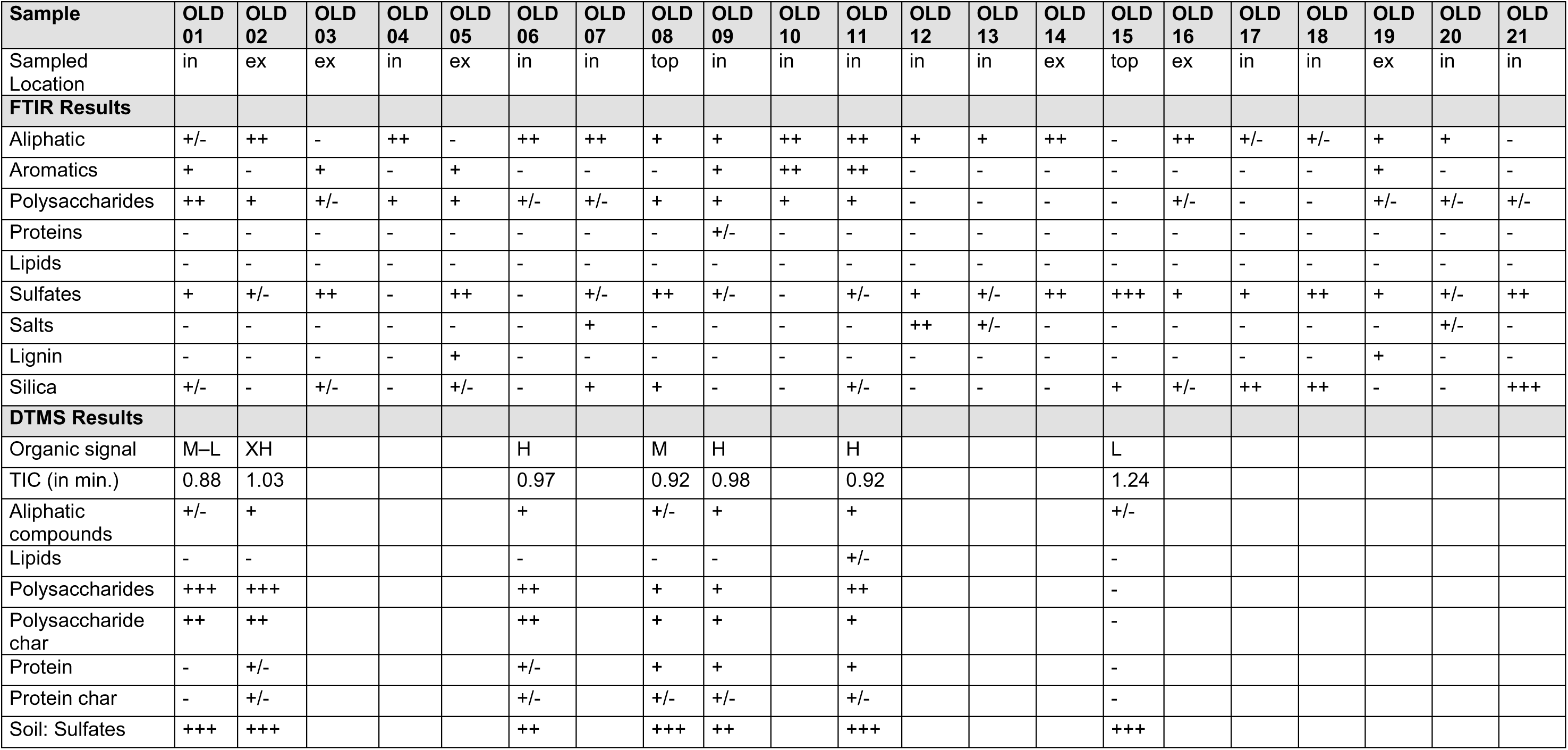
Summary of combined FTIR and DTMS results. The relative intensity of various characteristics is indicated in five categories (-: absent, +/-: minimal, +: average, ++: strong, and +++: very strong). Organic signal = cumulative intensity of the measurement, where XH = extra High (>300 x blank measurement), H = high (300–100 x), M = medium (100–40 x), L/M = low to medium (40–20 x) and L = low (<20x blank measurement)

Information about the original vessel contents could be obtained from 7 of the 21 residues (33%). In six cases, these residues were situated on the interior vessel wall and in one case, on the exterior vessel wall. Most prominent are the indications for the presence of well-preserved or slightly charred polysaccharides and the lack of significant amounts of lipids or proteins. While some residues contain minor amounts of proteins (in varying degrees of thermal degradation), polysaccharides form the main surviving class of compounds.

These 7 well-preserved residues can be grouped according to decreasing organic content and increasing thermal exposure. Three residues (OLD 02, situated on an exterior surface, and OLD 04 and 06, situated on interior surfaces) have high organic content and clearly show the presence of polysaccharides, identifiable by three peaks between 1000 and 1100 cm^-1^ (i.e., 1017, 1043, and 1080 cm^-1^) and in a single peak at 1151 cm^-1^. Significant aliphatic peaks at 2930 and 2850 cm^-1^, typical for aliphatic hydrocarbons in polysaccharides, are also present. The spectra show no characteristics of lipids or proteins; moreover, the lack of aromatic compounds indicates that the residues were barely carbonised (or charred). Three other residues (OLD 09, 10 and 11, all from the interior surfaces of a vessel) show a lower organic content, primarily polysaccharides, and, in the case of OLD 09, a minor amount of proteins. Here, the polysaccharides are well preserved, but the aliphatic signals are reduced and some aromatic compounds occur, indicating that the residues in this group have been exposed to more thermal degradation. In OLD 01, originally adhering to the interior surface of a vessel, even fewer organics could be detected. Very few aliphatic elements and only traces of polysaccharides are visible. This sample is of an aromatic nature, showing that the degree of thermal degradation is higher than in the other food crusts (Fig 18).

The presence of well-preserved polysaccharides and the absence of any significant amount of lipids or proteins is remarkable. When mixed foods are exposed to heat (e.g., during cooking), it is the polysaccharides that are the first to lose their unique characteristics. This is due to them being the most susceptible to thermal degradation, followed by proteins and then lipids.

In 8 of the 21 residues (38%), the FTIR spectra showed a good resolution, indicating that these samples contained enough material for an adequate ATR–FTIR analysis. Nevertheless, the analysis did not provide information about the original contents of the vessels. The residues (OLD 07, 08, 13, 14, 16 and 18–20) displayed low aliphatic signatures and a series of indications for various kinds of degradation and contamination from soil compounds. First, salts of carboxylic acids (identified by two wide bands, at 1595 cm^-1^ and 1395 cm^-1^) were found in some samples. These non-informative salts may originate from degraded lipids in the residue or the soil matrix. Second, some residues contain silicates (originating from sand or clay in the soil) and/or lignins (probably originating from modern roots or other plant material). The presence of these compounds indicates contamination from the soil substrate making these samples unsuitable for further analysis. Even though some of the residues in this group showed markers for polysaccharides (especially the residue on the fragment of disc OLD 08), identification of the original vessel contents was not possible due to the high degree of contamination (Fig 19).

**Fig 19.**
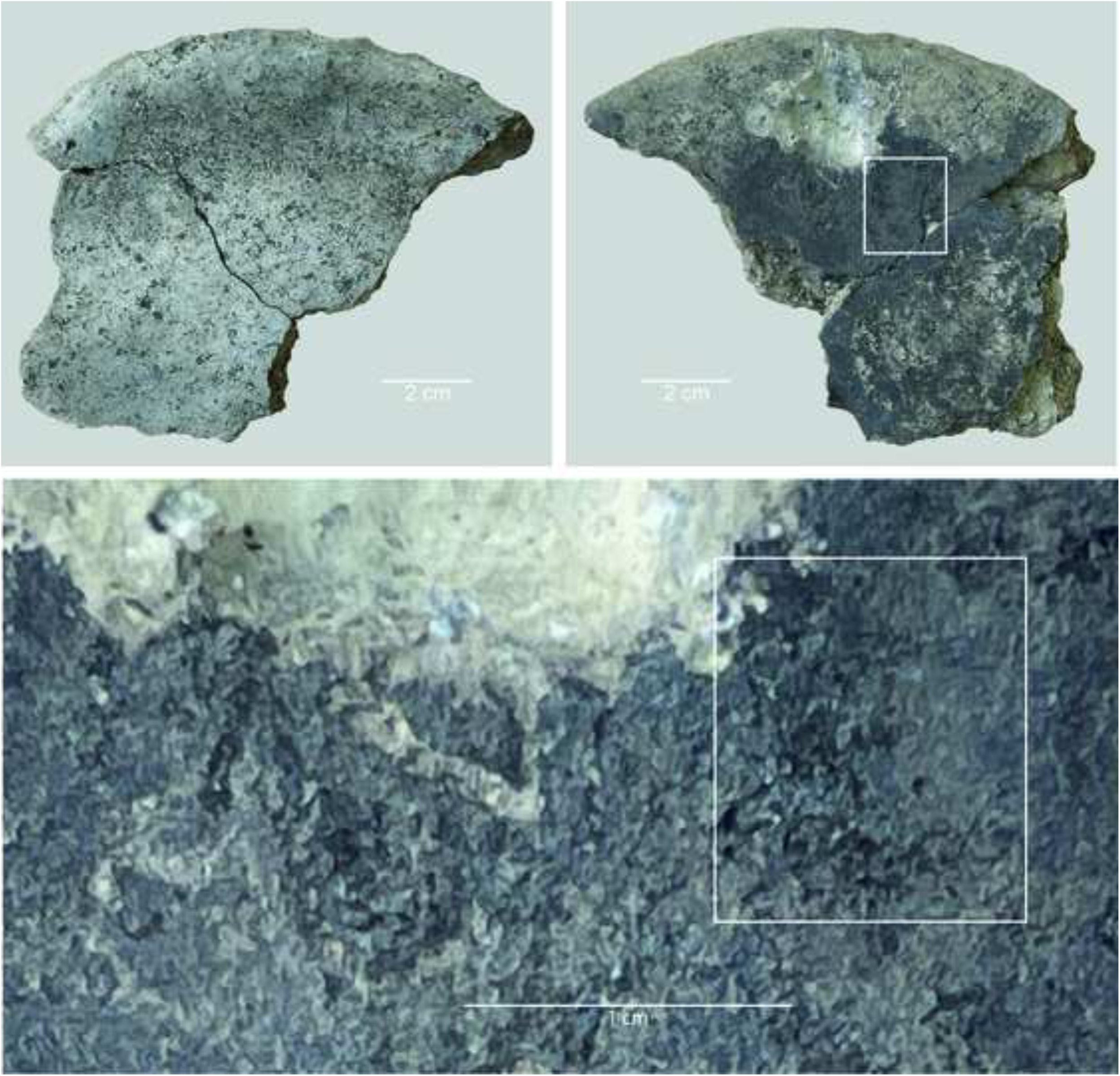
Oldenburg LA 77, sherd OLD 08: images show the bottom surface with signs of fire exposure (top left), and the top surface with residue (top right) as well as a detail of the sampled residue (bottom). White rectangles mark the sampled location. Images: T.F.M. Oudemans.

Of the 21 residues, 6 (29%) yielded no organic compounds; instead, their spectra exhibit the presence of sulfates. These 6 are OLD 03 and 05 (from the exterior surface); OLD 12, 17 and 21 (from the interior surface); and OLD 15 (from the surface of one of the ceramic discs) (Fig 20). Calcium sulfates are identified in the FTIR spectra by a combination of two wide bands, at 3526 and 3402 cm^-1^; a narrow band at 3242 cm^-1^; two narrow, sharp bands, at 1683 and 1620 cm^-1^; a broad band at 1445 cm^-1^; and a main band at 1108 cm^-1^ (see S4). Sulfates regularly occur in marine clay deposits. They can be formed by sulfur/sulfate-reducing bacteria, or due to the presence of sulfate-containing minerals (e.g. gypsum), or leaching at slightly acidic pH levels. They are present in most of the residues from Oldenburg LA 77 in varying amounts and can be considered contamination resulting from the local soil conditions.

**Fig 20.**
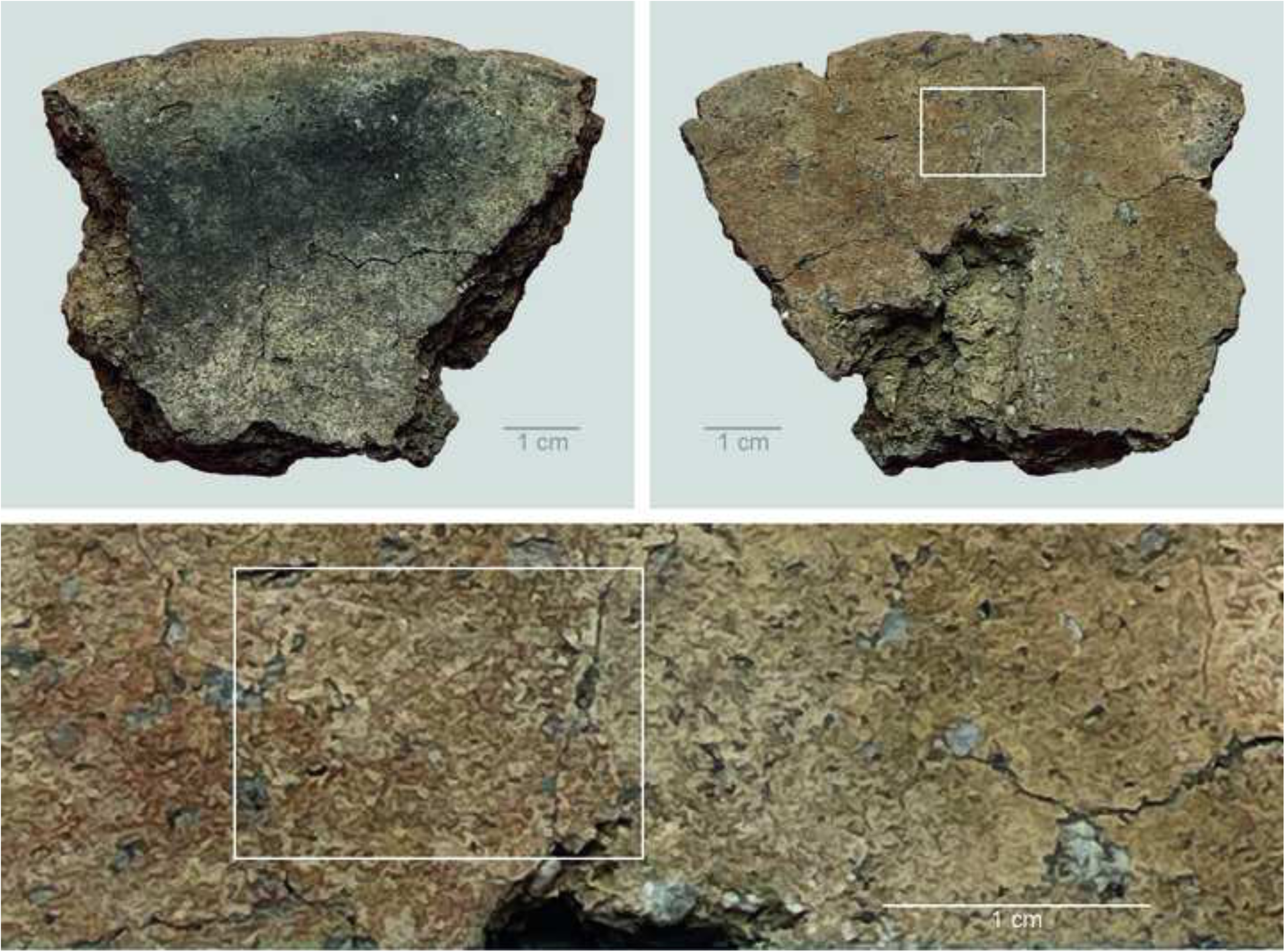
Oldenburg LA 77, sherd OLD 15: images show the bottom surface with signs of fire exposure (top left), and the top surface with residue (top right) as well as a detail of the sampled residue (bottom). White rectangles mark the sampled location. Images: T.F.M. Oudemans.

In summary, roughly one-third (33%) of the analysed residues contain well-preserved remains of the original foods cooked in the vessels, while two-thirds (67%) are clearly too contaminated with soil components to be indicative of the vessel contents. The 7 well-preserved residues primarily contain polysaccharides, demonstrating that the food consisted principally of starches and/or sugars. The origin of the polysaccharides cannot be determined in more detail, because the FTIR spectra of complex materials give only a general picture of the nature of the material. But since lipids are absent, a plant origin is considered the most likely. The composition of modern wholewheat grain shows similarities with some of the archaeological residues (S4). All well-preserved residues are situated on the surface of vessels: one is situated on the exterior of a vessel (14%) and the other six are internal residues on vessels (85%). Neither of the two residues from the ceramic discs displayed identifiable organic compounds in the FTIR analysis.

### DTMS results

A total of 7 residues, originally situated on the interior (n=4) or exterior (n=1) surface of vessels or the top surface of ceramic discs (n=2), were measured using DTMS (see Table 3). These residues exhibit different levels of thermal exposure. Even though the FTIR showed that the residues from ceramic discs are less promising for chemical analysis, they were included because the DTMS might help elucidate the function of these objects that has thus far remained unresolved. The results of each DTMS analysis form a series of mass spectra (see Fig 21), in which specific masses (or combinations of masses) represent known fragments of particular organic components or biomarkers in archaeological residues [38]. This makes it possible to score the presence or absence of various groups of organic molecules per residue (Table 3).

**Fig 21.**
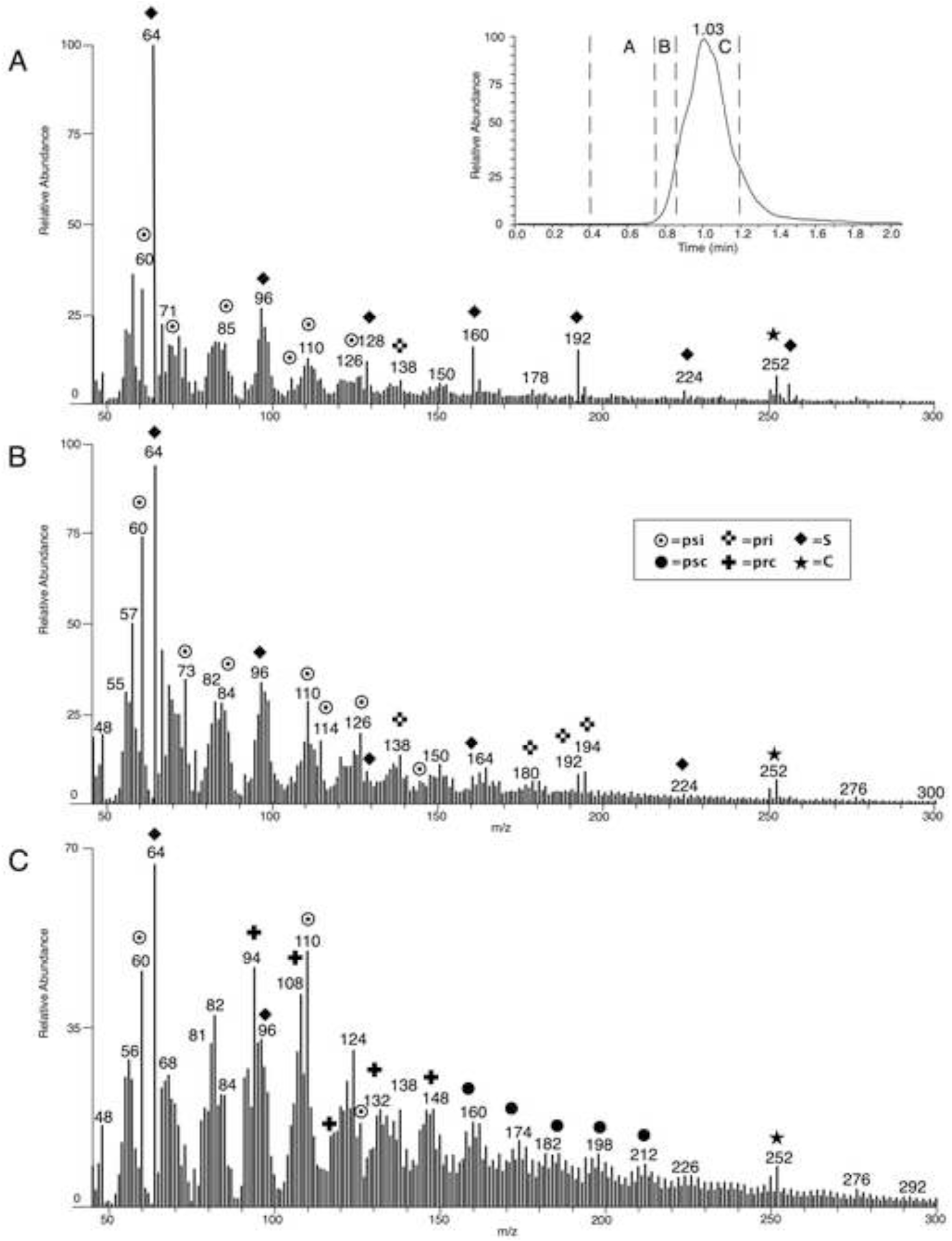
**DTMS results for residue OLD 02 including: the TIC or *total ion current* (top right) showing one peak at 1.03 minutes; the DTMS spectrum for the evaporation phase (area A, 0.4–0.76 minutes) (A); the DTMS spectrum for early pyrolysis phase (area B, 0.76–0.86 minutes) (B) and the DTMS spectrum for later pyrolysis phase (area C, 0.86– 1.20 minutes) (C)**. Markers for component classes are indicated with symbols: psi = intact polysaccharides; psc = thermally degraded and charred polysaccharides; pri = intact proteins; prc = charred proteins; S = elementary sulfur; C = contamination). Figure: T.F.M. Oudemans.

Six residues provided information on the original vessel contents; residue OLD 15, from one of the ceramic discs, contained too little organic material for a reliable interpretation as demonstrated by the cumulative intensity of the DTMS measurement, which is indicative of the organic content of the sample. For OLD 15 this response was only 9 times higher than the measurement without a sample where the common cut-off is at 20× the value. The DTMS results for the six samples confirm the findings of the FTIR analysis (Table 3): the samples contain polysaccharides, some better preserved than others. The DTMS analysis offered some additional information, in that it detected the presence of small amounts of partially charred proteins (in OLD 02, 08, 09 and 11) and a small amount of lipids in the form of free fatty acids (in OLD 11). Contamination with sulfur-containing compounds is also clearly visible in the DTMS analysis (see, for instance, the results for residue OLD 15). Further details on the DTMS results for the individual samples are given below.

Residue OLD 02 is rich in organic components and shows a high organic signal (Fig 21). The organic signal of OLD 02 is 547× higher than a measurement without sample. The total ion current (TIC) shows one, medium-wide, peak at an average temperature (maximum at 1.03 minutes), indicating a relatively homogeneous material of medium condensation, suggesting that some, but not much, thermal degradation had taken place (Fig 21). The desorption phase of OLD 02 (area A, 0.40–0.75 minutes) is dominated by markers for elementary sulfur (S2) that is liberated from the material during the measurement and originates from the sulfur-containing compounds in the sample. A very small amount of polysaccharides is also visible (Fig 21). Volatile components, such as lipids, normally visible in this part of the analysis, are not seen in OLD 02. As the temperature is increased during the early pyrolysis phase (area B, 0.76–0.85 minutes) a clear polysaccharide profile is visible, showing many intact polysaccharides as well as the remaining sulfur signal. Some markers for intact proteins are also visible. The later pyrolysis phase (area C, 0.87–1.20 minutes) shows primarily markers of thermally degraded polysaccharides, in combination with some minor compounds for charred proteins. Markers such as alkylphenols and alkylbenzenes (*m/z* 91, 92, 94, 105, 107, 108, 131) and nitrogen-containing compounds (*m/z* 117, 131, 147) indicate some thermal degradation of proteins. Residue OLD 02 consists mostly of polysaccharides and is a good example of the limited thermal degradation in the residues.

Polysaccharides usually originate from plant sources, but dairy products can also contain polysaccharides (in the form of milk sugars). However, since no lipids were detected, it can be concluded that the polysaccharides are of plant origin. The small amount of proteins visible in the sample are most likely part of the original, starch-rich plant material.

Residue OLD 01 is not very different from OLD 02, except for the fact that it contains less organic material. The TIC peak is much narrower and lies at lower temperatures, at 0.88 minutes, which indicates a very homogeneous material that is less condensed and thus less thermally degraded. The general composition is comparable to that of OLD 02, although it contains no protein fragments. However, due to the low organic content, it is possible that the protein fragments fall below the signal-to-noise ratio.

Residues OLD 06, 08 and 09 are similar to OLD 02 but contain less organic material. The narrow TIC peaks lie at lower temperatures, indicating relatively homogeneous materials of a less condensed nature and pointing to a lower degree of thermal degradation. The general composition is comparable to that of OLD 02, with some minor differences in the amount of (lightly charred) protein markers.

One of the residues from the ceramic discs (OLD 08) contained polysaccharides and no lipids, showing that the crust is probably of plant origin. Residue OLD 11 differs somewhat from the other residues, even though it contains a similar amount of organic material and the TIC peak lies around the same temperature (0.92 minutes). The general composition corresponds well to that of the other residues, but the sample contains a small amount of lipids, in the form of saturated free fatty acids. Although the lipid quantities are not significantly higher than the signal-to-noise ratio, minute traces of cholesterol indicate that a (partly) animal origin cannot be excluded.

In summary, all but one of the residues analysed with DTMS contained sufficient organic material for a reliable interpretation and thus yielded evidence of the original foods prepared or processed in the ceramic item. Of the 6 successful residues, 5 (83%) contain abundant polysaccharides and a minimal amount of proteins and lack lipids. The polysaccharides and proteins originate from starch-rich plant material. Only one residue (OLD 11) contained a small amount of lipids, which may indicate a (partly) animal origin. Five of the residues were found on the surface of a vessel (one situated on the exterior and the other situated on the interior of a vessel). One of the residues from a ceramic disc showed identifiable polysaccharides.

## Discussion

### Cooking with cereals

The results of the combined SEM, FTIR and DTMS analyses of surface residues from Oldenburg LA 77 (presented in Table 4) provide a significant contribution to the existing knowledge on prehistoric food processing and preparation techniques in northwestern Europe. Our study is the first of this kind both for northern Germany and for the Funnel Beaker groups.

**Table 4.** Overview of combined botanical and chemical results from Oldenburg LA 77 organic residues.

| Sample identifier | ALSH inventory number | Find number | SEM results | FTIR results<br>all samples contain sulfates unless otherwise indicated | DTMS results<br>all samples contain sulfates unless otherwise indicated | Interpretation of food crust |
| --- | --- | --- | --- | --- | --- | --- |
| OLD 01 | 2010-663.3082 | 232 | emmer grain fragments embedded in residue matrix, exposed aleurone & pericarp tissue | strong polysaccharide signal, minimal aliphatics, no proteins or lipids | medium to low in organics: mostly polysaccharides (intact and charred), no lipids or proteins | cooking/heating emmer grain, likely in water |
| OLD 02 | 2010-663.3087 | 309 | vegetative parenchyma, no further determination | strong polysaccharide signal, strong aliphatic signal, minimal proteins | very high in organics: mostly polysaccharides (intact and charred), minor amount of proteins (intact and charred), no lipids | cooking/heating starch- or sugar-rich plant material |
| OLD 03 | 2010-663.3096 | 423 | possible cereal grain endosperm matrix | polysaccharides, no aliphatics, no proteins or lipids, minimal silicate signal | *no DTMS* | possible cooking/heating cereal grain |
| OLD 04 | 2010-633.3099 | 154 | emmer grain fragments embedded in residue matrix, exposed aleurone & pericarp tissue | polysaccharides, strong aliphatic signal, no proteins or lipids | *no DTMS* | cooking/heating emmer grain, likely in water |
| OLD 05 | 2011-432.3093 | 382 | cf. bone fragment | strong polysaccharide signal, no aliphatics, minimal proteins | *no DTMS* | cooking/heating starch- or sugar-rich plant material |
| OLD 06 | 2010-633.3101 | 546 | emmer grain fragments, exposed aleurone & pericarp tissue; also seed testa of fat-hen (*Chenopodium album*) | strong polysaccharide signal, strong aliphatic signal, minimal proteins | high in organics: polysaccharides (intact and charred), no proteins or lipids | cooking/heating emmer grain with fat-hen seeds, likely in water |
| OLD 07 | 2010-633.3094 | 197 | *no SEM* | polysaccharides, strong aliphatic signal and silicates present | *no DTMS* | unknown residue with some contamination with soil material |
| OLD 08 | 2011-432.7915 | 2001 | *no SEM* | aliphatics, minimal polysaccharides, minimal proteins | medium in organics: polysaccharides (intact and charred), minor amount of proteins (intact and charred), no lipids | cooking/heating starch- or sugar-rich plant material |
| OLD 09 | 2011-432.7907 | 2158 | barley grains embedded in residue matrix | polysaccharides, strong aliphatic signal | high in organics: polysaccharides (intact and charred), minor amount of proteins (intact and charred), no lipids | cooking/heating unripe or milk-ripe barley grain, likely in water |
| OLD 10 | 2011-432.8108 | Ge-112 | barley grains embedded in residue matrix | minimal polysaccharides, strong aliphatic signal | *no DTMS* | cooking/heating unripe or milk-ripe barley grain, likely in water |
| OLD 11 | 2011-432.8096 | 2109-01 | emmer grain fragments embedded in residue matrix, exposed aleurone & pericarp tissue | polysaccharides, strong aliphatic signal, minimal proteins | high in organics: mostly intact polysaccharides (some charred), minor amount of proteins (intact and charred), some lipids | cooking/heating emmer grain, likely in water or milk |
| OLD 12 | 2011-432.8096 | 2109-02 | emmer grain fragments embedded in residue matrix, expose aleurone & pericarp tissue | no polysaccharides, some aliphatics, strong signal for salts of organic acids (degraded lipids) | *no DTMS* | strongly degraded residue of cooking/heating emmer grain, likely in water |
| OLD 13 | 2010-633.3073 | 5007 | *no SEM* | minimal polysaccharides and aliphatics, some salts of organic acids (degraded lipids) | *no DTMS* | unknown degraded residue |
| OLD 14 | 2010-633.3083 | 147 | *no SEM* | polysaccharides, strong aliphatic signal, minimal proteins | *no DTMS* | cooking/heating starch- or sugar-rich plant material |
| OLD 15 | 2010-633.3092 | 308 | *no SEM* | minimal polysaccharides, no aliphatics, strong sulfate signal, contamination with silicates | low in organics: minimal aliphatics, major sulfate contamination | soil material |
| OLD 16 | 2010-633.3100 | 334 | *no SEM* | minimal polysaccharides, strong aliphatic signal, no lipids or proteins | *no DTMS* | unknown residue |
| OLD 17 | 2010-633.3102 | 95 | *no SEM* | minimal polysaccharides, minimal aliphatics, minimal proteins, strong signal for salts of organic acids (degraded lipids) | *no DTMS* | unknown degraded residue |
| OLD 18 | 2011-432.7908 | 2361 | *no SEM* | minimal polysaccharides, minimal aliphatics, no lipids or proteins | *no DTMS* | unknown residue |
| OLD 19 | 2011-432.7908 | 2361 | *no SEM* | polysaccharides and aliphatics, no lipids or proteins | *no DTMS* | unknown residue |
| OLD 20 | 2011-432.7910 | 2440 | *no SEM* | minimal polysaccharides, aliphatics, strong signal for salts of organic acids (degraded lipids) | *no DTMS* | unknown degraded residue |
| OLD 21 | 2011-432.7912 | 2088 | *no SEM* | no polysaccharides, minimal aliphatics, significant silicate signal | *no DTMS* | soil material |

The picture emerging for Oldenburg LA 77 shows that cereals (specifically emmer and barley) formed the major ingredient of the dishes cooked in coarse-fabric vessels with thick walls. SEM analysis shows that these were apparently the only ingredients in all but one instance considered in this study. The exception is the combination of emmer and fat-hen seeds (in OLD 06). The addition of edible, protein-rich fat-hen seeds to the cereal-based meal is proposed to be deliberate rather than accidental. Further, the solid, or fused, appearance of the emmer grain endosperm in the Oldenburg LA 77 organic residues indicates that liquid, most likely water, was used in the cooking/heating process.

The FTIR analysis shows that one-third of the analysed surface residues contain traces of the original foods prepared or processed and that the residues are characterised primarily by polysaccharides (starches or sugars). The DTMS analysis confirmed the presence of relatively well-preserved polysaccharides but also provided the additional information, that most of the residues contain traces of proteins and that the residues contain no or only minute amounts of lipids (the latter in only one residue, OLD 11). The vast predominance of polysaccharides and the near absence of lipids make a plant origin the most likely for these residues. Whole cereal grains contain fair amounts (5–20%) of protein, and they are the most likely source of the proteins detected in the residues. The sample taken from the top surface of the fragment of one of the ceramic discs (OLD 08) revealed the presence of polysaccharides, suggesting that this object may indeed have been used to heat food containing cereal grains.

The presence of unripe barley grains in some of the residues is a peculiar aspect of Oldenburg food preparations. These would have been naked barley grains, as indicated by the analysis of archaeobotanical remains from the site. Ears of barley were probably harvested when they were still green. Due to the limited shelf life of green ears [42], they would need to have been processed and used soon after the harvest. The green barley harvest may have taken place in mid- or late summer (depending on whether sowing was done in winter or spring); assuming that the green barley was used soon after harvesting, these particular food remains may originate from late summer or early autumn cooking events.

The harvesting and consumption of green, unripe grains, primarily wheat and occasionally barley, are well documented ethnographically and described in publications by, for instance, [43-45]. However, there is only one published study that investigates different stages of maturity of cereal grains, more specifically spelt wheat grains (*Triticum spelta*), recovered from an archaeological site at Vaihingen in Germany, dating to the early La Tène period [42]. Therefore, the presence of immature barley grains in food crusts from Oldenburg represents direct evidence of a previously unrecorded culinary practice in Neolithic Europe.

The results presented here paint a different picture than that obtained from earlier studies of European Neolithic assemblages of food residues encrusted on pottery. The remarkably well-preserved food crusts on ceramic vessels from the Late Neolithic site of Pestenaker (3496–3410 BCE) in southern Bavaria contained evidence of the processing of mixed dishes and the cooking of combinations of cereals and dairy products [21]. A wide range of food sources (both plant and animal) were identified in mixed residues from the large ceramic assemblages from the Late Neolithic Corded Ware sites in the Netherlands (c. 2800–2400 BCE). At the site of Keinsmerbrug, for example, emmer was cooked with the addition of animal fat, while at the sites of Mienakker and Zeewijk, dishes comprised a mixture of foods [32-33, 46].

Consistent with an earlier study [47], we have proven that terrestrial foods were cooked in pots in this early farming village. However, the vast predominance of polysaccharides and the near absence of lipids from the surface residues stand in contradiction with the findings from the earlier work. Weber and colleagues extracted lipids from the pottery fabric of potsherds (n=18) from Oldenburg LA 77 and argued that the absorbed lipids represent the dishes cooked in these vessels. They found that some of the sherds contain indicators for food of plant origin (n=9), milk or dairy products (n=5), and food of meat origin (n=2); two sherds contained no significant amount of lipids. The discrepancy between this and the results of our study is significant: because lipids are much more heat-resistant and less soluble than polysaccharides, lipids would have been detected in the surface residues if they had been present in the dishes cooked in the vessels that we analysed. In both studies, a similar set of heavily fragmented and undecorated ceramic sherds were analysed, originating from vessels thought to have been in regular, everyday use [1-2]. The discrepancy could perhaps be explained by a possibility that the lipids discovered in the ceramic matrix were the result of a post-firing waterproofing technique, involving the use of milk, dairy products, or plant oil, intended to make the pottery less permeable—the practice documented ethnographically and successfully tested experimentally [48-49].

### Processing and preparation: sprouting and dehusking of emmer

The absence of emmer chaff from the emmer-containing food crusts from Oldenburg points to thorough processing—dehusking, sieving, and winnowing—prior to food preparation. The archaeobotanical assemblage includes abundant remains of emmer chaff and, thus, indirectly, evidence for emmer having been stored in spikelets and for the final stages of cleaning having been done on-site. Parching of spikelets is often cited as a method applied to ease the removal of chaff, but the presence of sprouted emmer grains in Oldenburg LA 77 food crusts and the lack of chaff in food preparations may reflect an alternative or additional strategy, such as soaking emmer spikelets before dehusking.

Dampening of spikelets softens the chaff and, during pounding, facilitates the release of the grain and removal (literally shredding) of the chaff through rubbing of the spikelets against each other and the dehusking device, without causing (excessive) damage to the grains. Experimental work and archaeobotanical evidence suggest that mortars and pestles could have been used in this procedure [50-51]. Soaking in water before dehusking, followed by either sun-drying or artificial drying or roasting in ovens, is also attested ethnographically [52] and references therein]. The result is a damp mass consisting of shredded fine and coarse glume wheat chaff and clean grain, which can be separated by sieving. Long periods of dampening, however, can lead to sprouting of the grain, and this scenario seems to have played out at Oldenburg. Moreover, sprouting seems to have been a desired outcome, given that the dampening occurred prior to dehusking—the logic being that pounding or grinding of dry emmer spikelets, which are hard and have chaff tightly enclosing the grain, would have damaged the embryo, which was needed for germination, however, in this case, the embryo was clearly intact enough to have been able to sprout. Soaking emmer spikelets would thus have served a dual purpose, namely, to facilitate the dehusking process and to induce germination. Analysis of the morphology of archaeological sprouted emmer grains from the New Kingdom site at Amarna in Egypt has shown that the chaff had been in place during the germination [50-51].

In Oldenburg LA 77, the soaking of spikelets likely occurred in a controlled environment, such as ceramic vessels, in low-light conditions. The soaking water would have been drained off after some time and the spikelets left wet. If the length of sprouts observed on some germinated ancient grains can be used as an indicator, then emmer grains may have sprouted within six to eight days [50-53]. Once free of chaff, the sprouted emmer grains at Oldenburg LA 77 would have been processed further before cooking, possibly by being crushed or ground, which would account for the presence of grain fragments embedded in otherwise featureless residue matrices in some of the food residues (OLD 01, 04, 11 and 12).

Sprouting is a natural biochemical process activated when seeds and grains are exposed to moisture and warmth, and sprouted grain has several advantages over unsprouted grain for humans. It triggers enzymes to break down stored nutrients—complex carbohydrates into simple sugars, proteins into amino acids, and fats into fatty acids—making the grains easier to digest. Sprouted grains may also have higher levels of vitamins and minerals, including vitamin C and iron. Overall, sprouting significantly increases the nutritional value of the grains, which is crucial for the growth of a new plant and beneficial for humans [54-55].

Sprouting also enhances the flavour of the grain. Foods prepared from sprouted grains have a distinctly sweet and nutty flavour compared with those made from unsprouted grains, most likely due to the conversion of starches into simpler sugars during the sprouting process. Finally, grain germination is a key step in the preparation of malt, which is an essential ingredient in brewing beer.

The Neolithic inhabitants of Oldenburg LA 77 may have recognised some of the advantages of inducing germination of emmer grains and the consumption of such grains, perhaps in the form of porridge or a malt-containing drink. Maybe some of the encrusted residues represent emmer mash intended for the production of fermented drinks. The generally low degree of thermal degradation of the organic materials in several of the Oldenburg LA77 residues, as indicated by the DTMS results, may signal exposure to low heating temperatures, which would be a desirable quality in the production of malted drinks.

The use of sprouted cereal grains has been reported for Late Neolithic lakeshore settlements in central Europe, including Hornstaad–Hörnle IA and Sipplingen–Osthafen, on Lake Constance, in southern Germany, and Zürich–Parkhaus Opéra, on Lake Zürich, in Switzerland. At all three settlements, sprouted grains were subjected to cooking or heating, resulting in either a malt-containing liquid (Hornstaad–Hörnle IA) or an ingredient that was in either liquid or solid food that contained malt (Sipplingen–Osthafen and Zürich–Parkhaus Opéra) [34]. These findings demonstrate that the processing of malted grain was practiced as part of various food production sequences during the 4th millennium BCE in central Europe.

## Conclusion

The combination of complementary high-resolution techniques—SEM analysis of the morphology and anatomy of plant remains and ATR-FTIR and DTMS for the identification of organic components—not only identified remains of food in surface residues on ceramics from the Funnel Beaker Neolithic in northern Germany, but also revealed some previously unknown aspects of food processing and preparation, and their timing, in the prehistory of northwestern Europe.

SEM study of the residues provided detailed insight into the ingredients and their state, identifying previously unknown ways in which cereal-based foods were processed and prepared for consumption, including the use of unripe barley grains, sprouted emmer grains, and a combination of cereals and wild plants in cooked meals. The chemical analyses demonstrated the presence of polysaccharides in combination with small amounts of proteins, and the absence of lipids indicating starch- or sugar-rich plant-derived foods as the main constituent of the residues. The abundance and chemical preservation of the polysaccharides indicates that these foods have only been heated at low temperatures.

The entire *chaîne opératoire* associated with Neolithic cereal production and consumption at Oldenburg LA 77 can now be traced—from managing the arable field, harvesting, and preparing the grain for storage, to preparing food in cooking pots or on baking plates. The time around 3100 BCE was the time when agricultural production in the region was being consolidated and people were beginning to group in villages, here exemplified by the Funnel Beaker community of the settlement at Oldenburg LA77. The inhabitants directed great attention to the plant food economy, not only in working and maintaining the arable land and tending to staple crops but also in processing and cooking the resulting grain.

Earlier studies of archaeological food remains had already demonstrated that prehistoric food and drink preparation and cooking were as complex and diverse as the evidence they left behind. This work has expanded the existing knowledge geographically and temporally and demonstrated the research potential of the multi-method approach. Most significantly, it has enabled us to get a glimpse of the final stages of the long process of transformation of food plants, from sowing to consumption.

## Acknowledgments

LK-M expresses her gratitude to the Naturalis Biodiversity Center in Leiden, the Netherlands, where she carried out the scanning electron microscope analysis using the JEOL JSM-6480L device. The ATR–FTIR measurements were performed at the Kenaz Consult & Laboratory in Berlin. We are indebted to Ester S.B. Ferreira (Cologne Institute of Conservation Sciences, University of Applied Sciences in Cologne) for providing DTMS accessibility; Andreas G. Heiss (Österreichische Akademie der Wissenschaften, Österreichisches Archäologisches Institut, Vienna) for his valuable thoughts concerning the taxonomic determination of immature barley grains; and Wouter van der Meer (BIAX Consult, Zaandam, the Netherlands) for measuring the thickness of the aleurone cell walls in emmer remains with the use of R-STUDIO software and producing the diagrams. We thank Agnes Heitmann and Karin Winter (Graphics Department of the Institute for Pre- and Protohistory in Kiel) and Daniel Knitter (OEKO-LOG field research) for their help with photography and illustration, and Suzanne Needs-Howarth (Toronto) for copy-editing. We are grateful to the editors and reviewers for the careful reading of our manuscript and useful suggestions.

## Credit Statement

L.K-M: Microscopic and SEM analyses, photography, characterisation and identification of the remains, data curation and interpretation, writing and editing. T.O.: photography, chemical analyses, characterisation and identification of the remains, data curation and interpretation, writing and editing. J-P.B.: archaeological excavation, data curation, writing and editing. D.F.: data curation, writing and editing. J.M.: archaeological excavation, data curation, funding acquisition. W.K. research coordination, writing and editing, funding acquisition.

## Supporting information

S1 Fig. Examples of thick-walled, undecorated pottery sherds from Oldenburg LA 77

S2 Fig and S2 Table. Radiocarbon AMS dates and their calibrated values for three samples of food crusts (OLD 04, 06, and 11) and two charred emmer grains (SED 27 and 28) (S2 by D. Filipović)

S3 Table 1. Aleurone double cell wall thickness measurements on tangential (top view)

S3 Table 2. Aleurone double cell wall thickness measurements on transverse (side view)

S4 Table. ATR–FTIR results. The relative intensity of the FTIR transmission bands is indicated in five categories (-: absent, +/-: minimal, +: average, ++: strong and +++: very strong). The origin of the bands is indicated as Mi = minerals; PS = polysaccharides; Pr = proteins; Al = aliphatic hydrocarbons; Ar = aromatic compounds; Lig = lignins; S = sulfates and Si = silicates. The type of transmission band is given as: s. = stretch; b. = bend; vibr. = vibration [such as rocking, scissoring, wagging, or twisting]; def. = deformation; skel. = skeletal; asym. = asymmetrical.

